# Robust virtual staining of landmark organelles

**DOI:** 10.1101/2024.05.31.596901

**Authors:** Ziwen Liu, Eduardo Hirata-Miyasaki, Soorya Pradeep, Johanna Rahm, Christian Foley, Talon Chandler, Ivan Ivanov, Hunter Woosley, Tiger Lao, Akilandeswari Balasubramanian, Rita Marreiros, Chad Liu, Manu Leonetti, Ranen Aviner, Carolina Arias, Adrian Jacobo, Shalin B. Mehta

## Abstract

Correlative dynamic imaging of cellular landmarks, such as nuclei and nucleoli, cell membranes, nuclear envelope and lipid droplets is critical for systems cell biology and drug discovery, but challenging to achieve with molecular labels. Virtual staining of label-free images with deep neural networks is an emerging solution for correlative dynamic imaging. Multiplexed imaging of cellular landmarks from scattered light and subsequent demultiplexing with virtual staining leaves the light spectrum for imaging additional molecular reporters, photomanipulation, or other tasks. Current approaches for virtual staining of landmark organelles are fragile in the presence of nuisance variations in imaging, culture conditions, and cell types. We report training protocols for virtual staining of nuclei and membranes robust to variations in imaging parameters, cell states, and cell types. We describe a flexible and scalable convolutional architecture, UNeXt2, for supervised training and self-supervised pre-training. The strategies we report here enable robust virtual staining of nuclei and cell membranes in multiple cell types, including human cell lines, neuromasts of zebrafish and stem cell (iPSC)-derived neurons, across a range of imaging conditions. We assess the models by comparing the intensity, segmentations, and application-specific measurements obtained from virtually stained and experimentally stained nuclei and cell membranes. The models rescue missing labels, non-uniform expression of labels, and photobleaching. We share three pre-trained models (VSCyto3D, VSNeuromast, and VSCyto2D) and a PyTorch-based pipeline (VisCy) for training, inference, and deployment that leverages current community standards for image data and metadata.

## Introduction

Building predictive models of complex biological systems requires correlative analysis of the dynamics of cells and organelles (1–6). Genetic tagging with multiple fluorescent proteins is a current gold standard for multiplexed imaging of organelle dynamics(7, 8). Despite advances in gene editing technologies fueled by CRISPR, labeling multiple organelles with fluorescent proteins is labor-intensive, particularly in cell models that recapitulate the native processes of an organism, such as development and neuronal differentiation. For example, analyzing the emergence and homeostasis of cell types during the development of the zebrafish neuromast (9, 10), requires tracking individual cell types and developmental signals. However, engineering embryos to express fluorescent reporters for developmental signaling, cell type, nuclei, and membranes is time-consuming. Fluorescent tags themselves, as well as phototoxicity caused by imaging multiple fluorescent channels, often compromise cell health. These trade-offs are compounded in high-throughput experiments that incorporate diverse perturbations and cell types. Multiplexed imaging of organelles and cell membranes over time and across perturbations is slowed down by sequential multispectral fluorescence imaging. In all of these cases, photobleaching of fluorophores limits the temporal resolution and the duration of experiments.

Virtual staining of quantitative label-free imaging data is a promising and pragmatic solution to the challenges mentioned above. 3D quantitative phase imaging methods (11–16) consistently visualize several landmark organelles - including nuclei, cell membrane, nucleoli, nuclear envelope, and lipid droplets - in a single image by measuring their dry mass. Quantitative polarization imaging methods measure the alignment and orientation of ordered organelles such as cytoskeleton, and can be multiplexed with quantitative phase imaging (11, 13, 17, 18). Raman microscopy also reports several organelles based on relative concentrations of nucleic acids, amino acids, and lipids (19, 20). If the distribution of these physical and chemical properties is correlated with the distribution of fluorescent labels, deep learning models can demultiplex organelles observed simultaneously by label-free contrast (11, 18, 21–24). Virtual staining also circumvents the need for laborious and error-prone human annotations of organelles in 3D volumes and movies. Beyond the analysis of cell dynamics, virtual staining is now widely used for rapid 3D histology from autofluorescence, optical coherence tomography, and Raman microscopy (25–28). If the organelles, cells, or tissue architecture of interest are consistently encoded by label-free contrast, virtual stains are more reproducible than experimental stains that rely on genetic engineering or tissue fixation (18) and chemical stains (27).

Above work on virtual staining suggests that it can indeed relax the longstanding multiplexing bottleneck in dynamic imaging. Then, why is this approach not used more widely? One of the outstanding challenges (24, 29) is that current virtual staining models, like most deep neural networks, do not generalize to imaging parameters, cell states, and cell types beyond the distribution represented by the training data. In this paper, we tackle this challenge by reporting strategies to train generalist virtual staining models that jointly predict nuclei and cell bodies across imaging conditions, cell states, and cell types.

Segmentation of nuclei and cells is a common first step in image-based phenotyping, given the heterogeneity of cellular responses. Several generalist segmentation models have been reported with diverse architectures (30–33) to segment nuclei and cells from fluorescence images. The majority of these models were not trained with high-resolution quantitative label-free imaging data and do not generalize to these datasets. Fine-tuning these models for quantitative label-free images requires expensive human annotation. We show that the combination of generalist virtual staining with off-the-shelf generalist fluorescence segmentation models enables reliable single-cell analysis.

This paper makes the following specific contributions: a) physics-based deconvolution and data augmentation strategies that make the virtual staining models invariant to nuisance changes in imaging parameters, and generalize them to a different phase contrast without requiring experimental training data, b) training protocols that generalize virtual staining models to cell states observed during development and differentiation, c) training protocols that generalize the virtual staining models to multiple cell types with minimal training data, d) a scalable convolutional image translation architecture (UNeXt2), and e) trained models for virtual staining of nuclei and membrane from widely deployable Zernike phase contrast or quantitative phase contrast data. Additionally, we provide a permissively licensed pythonic pipeline, named <u>VisCy</u> (34), for model training, inference, and evaluation that implements these strategies. We assess the gains in performance due to architectural refinement, augmentation strategies, and training protocols using a suite of metrics that include regression metrics, instance segmentation metrics, and application-specific metrics.

## Results

### Data, architecture, and metrics

Virtual staining models are trained using paired label-free and fluorescence images (Figure 1A, orange and blue arrows). Once a model is trained, only the label-free input is needed for inference (Figure 1A, orange arrows). We focus on training the models with an inexpensive quantitative phase imaging (QPI) method of “phase from defocus” (11, 35, 36), which can be implemented on any motorized widefield microscope. This straightforward QPI method consists of acquiring a z-stack in transmission and deconvolving phase density (<u>Methods</u>-<u>Preprocessing</u>) using an optical transfer function that maps the specimen’s 3D phase density (i.e., local dry mass) to the 3D contrast variations.

**Figure 1.**
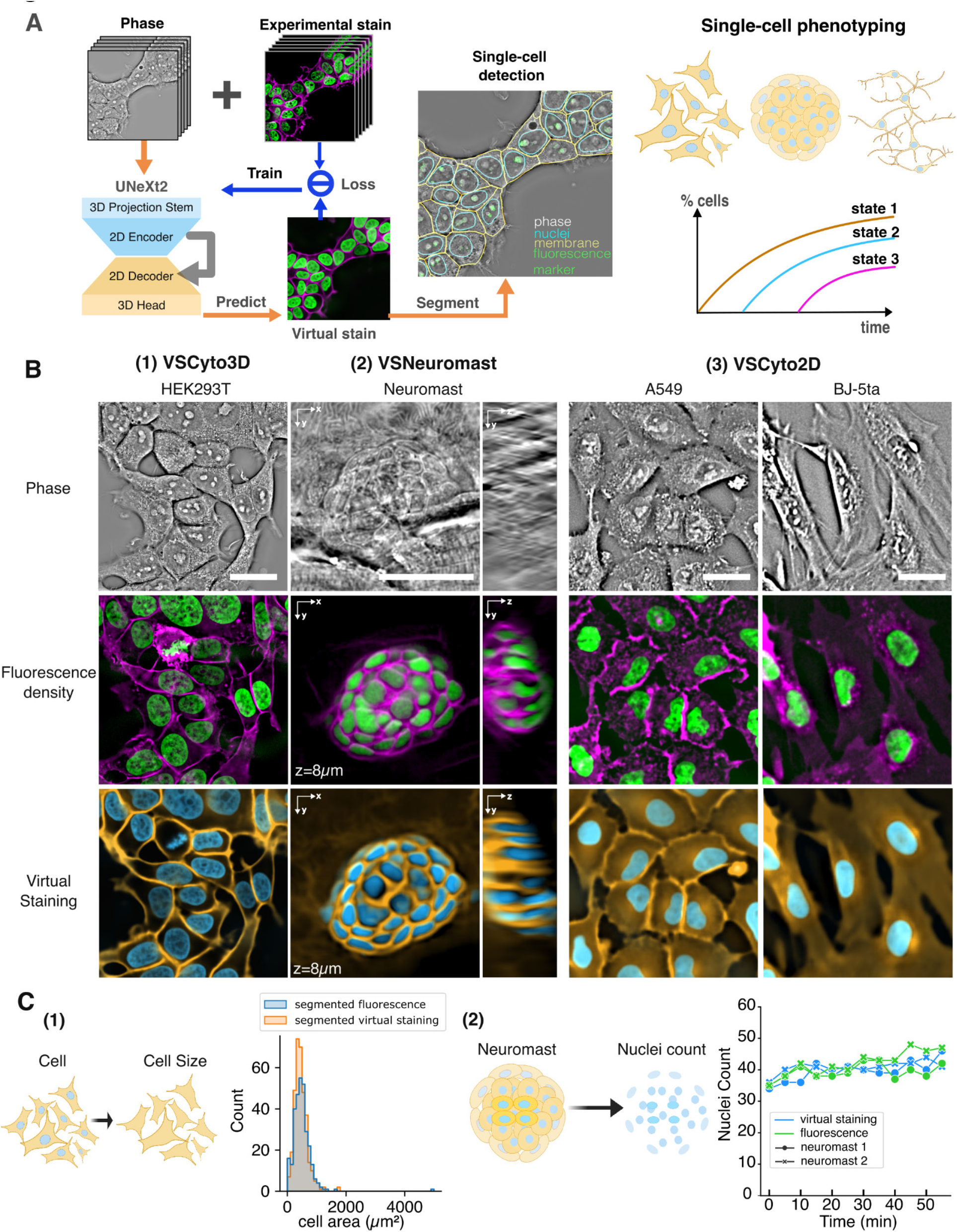
Robust virtual staining **(A)** Schematic illustrating the training (blue arrows) and inference (orange arrows) processes of robust virtual staining models using UNeXt2 with physically informed data augmentations to enhance performance and generalizability. The models virtually stain nuclei and membranes, allowing for single-cell phenotyping without experimental staining. We utilize generalist segmentation models to segment virtually stained nuclei and membranes. **(B)** Input phase images (top row), experimental fluorescence images of nuclei and membrane (middle row), and virtually stained nuclei and membrane (bottom row) using VSCyto3D (HEK293T cells), fine-tuned VSNeuromast (zebrafish neuromasts), and VSCyto2D (A549 and BJ-5ta cells). Virtually and experimentally stained nuclei and membranes are segmented using the same Cellpose model. The instance segmentations are compared using the average precision at IoU of 0.5 (AP@0.5). Scale bars: 25 µm. **(C)** We rank and refine models based on application-specific metrics, in addition to instance segmentation metrics. (1) Morphological Measurements: We compare cell area in HEK293T cells measured with segmentation of experimentally and virtually stained membranes. (2) Nuclei Count: We compare the number of nuclei in neuromasts identified from experimentally and virtually stained nuclei over short and long developmental time windows. The plot shows the number of nuclei over one hour, measured every 5 minutes on 3 dpf fish.

With the primary goal of accelerating single-cell phenotyping, we developed models for joint virtual staining of nuclei and cell membranes (Figure 1B) that address distinct use cases: 3D virtual staining of HEK293T from the OpenCell library of endogenously-tagged human cells lines (VSCyto3D), 3D virtual staining of zebrafish neuromasts for analyzing cell growth and death during development (VSNeuromast), and 2D virtual staining for high throughput screens across multiple cell types, HEK293T, A549, BJ-5ta, and induced neurons (iNeurons) (VSCyto2D). In all of these applications, virtual staining and generalist segmentation models are used in tandem to segment the nuclei and cells from label-free images. Combination of QPI and complementary fluorescence sensors then enable phenotyping of functional states with single-cell resolution (Figure 1A, single-cell phenotyping).

We use a *purely convolutional* architecture that draws on the design principles of transformer models. A variety of convolution and attention-based architectures have been reported for image translation and cell segmentation. There is an active debate whether transformer models that use attention operation (31–33, 37) fundamentally outperform convolutional neural networks that rely on the inductive bias of shift equivariance. Systematic comparisons suggest that convolutional models perform as well as transformer models (37, 38) when large compute budget is spent, and outperform the transformer models when moderate compute budget is spent. Therefore, we integrate the design choices from UNet (39), convNeXt v2 (40), and SparK (41) architectures to develop an efficient and scalable architecture, named UNeXt2 (Figure 1A).

UNeXt2 architecture can be used for 2D, 3D, or 2.5D (11) image translation. The module in the network that enables flexible choice of number of slices in the input stacks and output stacks is a projection module in the stem and head of the network. The body of the network is a UNet-like hierarchical encoder and decoder with skip connections that learns a high-resolution mapping between input and output. The choice of layers, blocks, and the loss function are described in <u>Methods-Model</u> <u>architecture</u> and Table 2. The UNeXt2 architecture provides 15x more learnable parameters for 3D image translation than our previously published 2.5D UNet at the same computational cost (Table 1). The efficiency gains are even more significant when compared to 3D UNet. This approach enables the allocation of the available computing budget to train moderate-sized models faster or to train more expressive models that generalize to new imaging conditions and cell types. Interestingly, the models trained for joint prediction of nuclei and membrane are slightly more accurate than models trained for prediction of nuclei alone (compare metrics shown in Table 1).

**Table 1.**
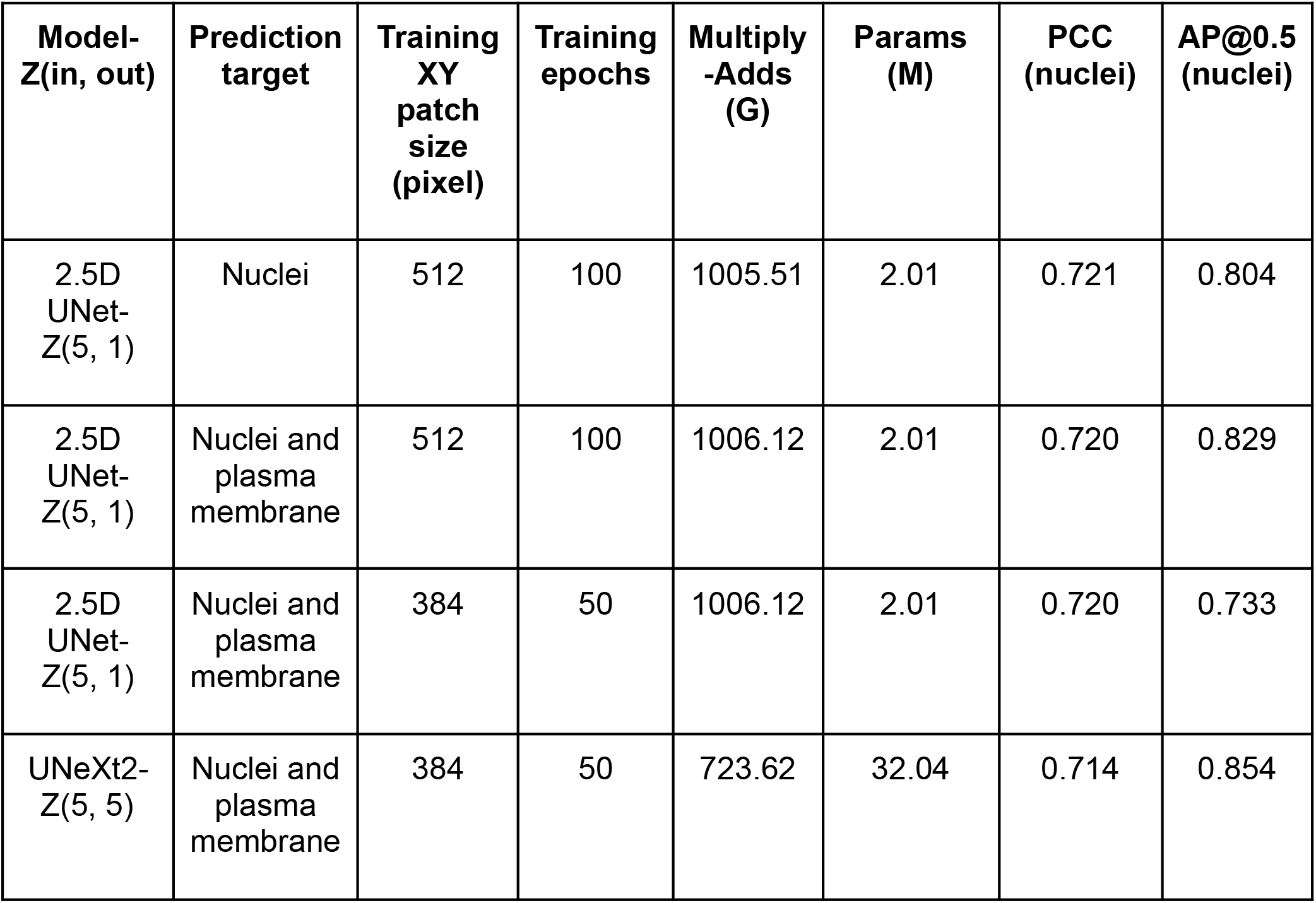
Comparison of the computational complexity, capacity, and performance of UNeXt2 models with previously published 2.5D UNet model: We compare metrics of accuracy of regression (Pearson correlation coefficient, PCC) and instance segmentation (average precision at IoU threshold of 0.5, AP@0.5) of nuclei on the central slice of HEK293T images. The computational complexity is measured with the number of multiply-add operations during inference on ZYX input size (5, 2048, 2048) with batch size 1. The model capacity is measured with the number of learnable parameters. Predicting both nuclei and membrane targets improves nuclei prediction with 2.5D UNet. UNeXt2 architecture provides higher learning capacity than 2.5D UNet architecture at similar computational complexity. Predictions with UNeXt2-Z(5,5) are shown in Figure 1.

| Model-Z(in, out) | Prediction target | Training XY patch size (pixel) | Training epochs | Multiply-Adds (G) | Params (M) | PCC (nuclei) | AP@0.5 (nuclei) |
| --- | --- | --- | --- | --- | --- | --- | --- |
| 2.5D UNet-Z(5, 1) | Nuclei | 512 | 100 | 1005.51 | 2.01 | 0.721 | 0.804 |
| 2.5D UNet-Z(5, 1) | Nuclei and plasma membrane | 512 | 100 | 1006.12 | 2.01 | 0.720 | 0.829 |
| 2.5D UNet-Z(5, 1) | Nuclei and plasma membrane | 384 | 50 | 1006.12 | 2.01 | 0.720 | 0.733 |
| UNeXt2-Z(5, 5) | Nuclei and plasma membrane | 384 | 50 | 723.62 | 32.04 | 0.714 | 0.854 |

**Table 2.**
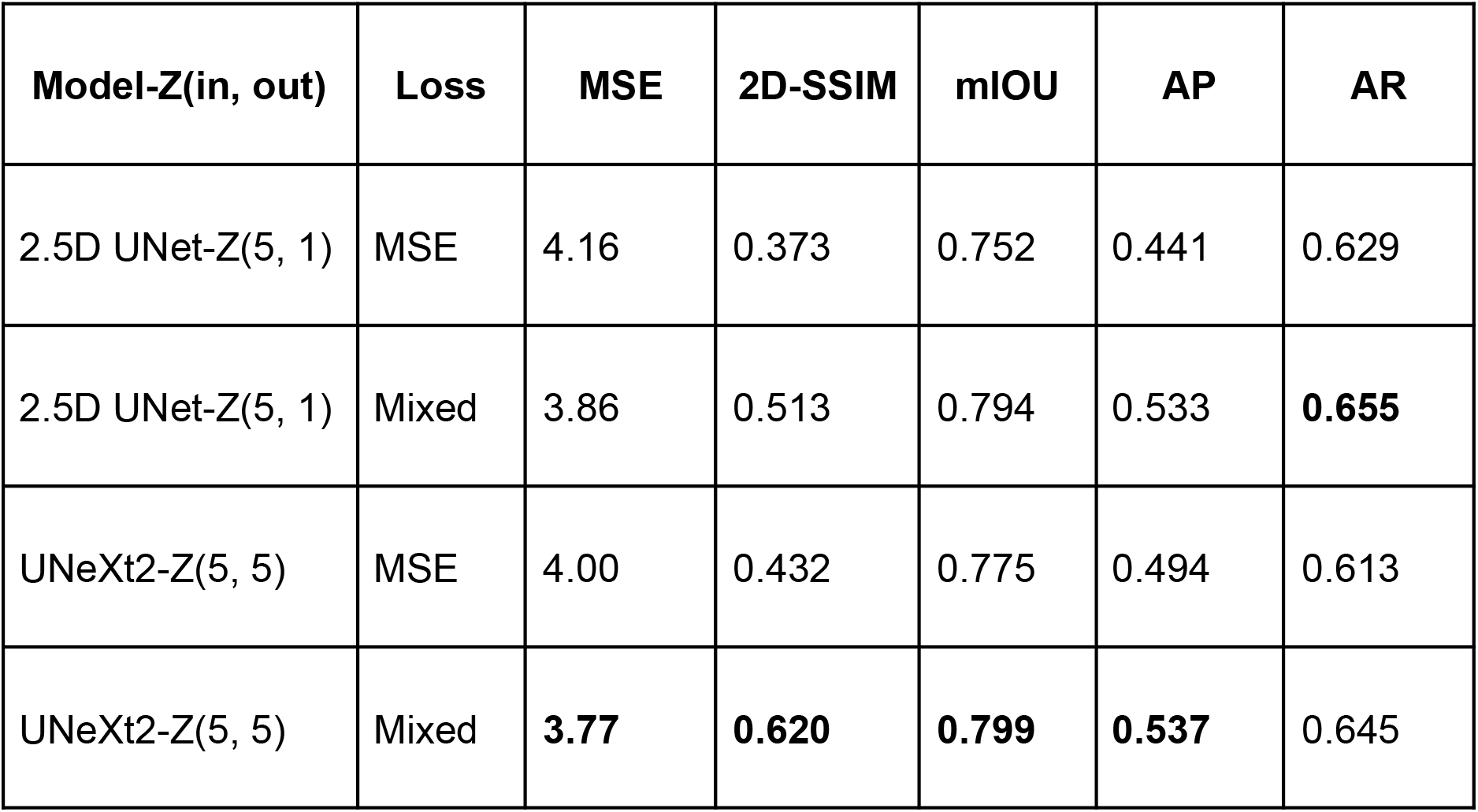
Comparison of the performance of models trained with 2 loss functions on the center slice of the HEK293T test dataset. The models trained with the mixed loss also have lower test MSE than models trained with the MSE loss

We use Cellpose(30) (see <u>Methods-Model Evaluation</u>, <u>Figure S1</u>) for segmenting the virtually stained nuclei and membrane. Joint virtual staining of nuclei and membranes enables more accurate cell segmentation (30). The Cellpose model requires significant fine-tuning with QPI images but works well with virtually stained images of nuclei and cell membrane, primarily because the training set of Cellpose included only classical Zernike phase contrast (42) and fluorescence data. As seen in Figure 1B, virtually stained images are intrinsically denoised because the models cannot learn to predict random noise. This feature obviates the need to train models that are robust to noise, such as Cellpose3 (31).

We assess the performance of the models using regression metrics (Pearson correlation), instance segmentation metrics (average precision), and application-specific metrics (e.g., cell count, cell area). Due to the variations in experimental labeling and the need to fine-tune Cellpose models to new cell shapes, we cannot rely on experimental fluorescence images and segmentations obtained with Cellpose as absolute ground truth. For example, boundaries of BJ-5ta cells at low magnifications (Figure 1B, <u>Figure S1</u>) are challenging to segment, because they have diverse shapes and grow on top of each other. Therefore, this paper first compares the experimental and virtually stained images and their segmentations, followed by quantifying the observations with metrics. The instance segmentations are compared using average precision (AP) between segmented nuclei (or cell membranes) from fluorescence density images and from virtually stained images. An instance of a cell is considered true positive if the intersection over union (IoU) of both segmentations reaches a threshold. We compute AP at IoU of 0.5 (AP@0.5) to evaluate correspondence between instance segmentations at the coarse spatial scale and mean AP (mAP) across IoU of 0.5-0.95 to evaluate the correspondence between instance segmentations at the finer spatial scales. Model refinement and hyperparameter optimization are guided by application-driven metrics such as cell size of cultured cells and nuclei count in neuromasts (Figure 1C), in addition to regression and segmentation metrics.

VSCyto3D and VSNeuromast are trained in a supervised fashion using UNeXt2 architecture and mixed loss. VSNeuromast is fine-tuned to new developmental stages using sparsely sampled time series of label-free and fluorescence volumes. VSCyto2D model is trained with a pre-training/fine-tuning paradigm common for language and vision transformers. Subsequent results describe each of these training protocols and our findings on the regime of generalization of the resulting models.

### Robustness to imaging parameters

Nuisance variations in label-free images, such as change in the type of phase contrast and optical aberrations, degrade the performance of virtual staining models. Common strategies to improve the robustness of the models is to include more training data, either by acquiring data under diverse conditions and by synthesizing training data by augmentation of the existing training data. Generating well-registered experimental training data for image translation across a variety of imaging conditions is onerous. Since the image formation process of microscopes can be modeled accurately, we reasoned that deconvolution of raw data and the augmentation of deconvolved data informed by the image formation process of microscopes can lead to virtual staining models robust to variations in imaging parameters. These computational experiments led to the VSCyto3D model.

The effect of deconvolution on performance of virtual staining models is evaluated by training four virtual staining models that translate between combinations of raw and deconvolved images using UNeXt2 architecture (<u>Figure S2</u>) as shown in Figure 2A. Deconvolution of raw intensities at the preprocessing stage (<u>Methods</u>-<u>Preprocessing</u>) can remove nuisance variations due to imperfect imaging conditions and improve contrast of biological structures in the image data. Deconvolution removes slowly varying intensity and phase variations that typically arise due to non-uniform illumination or meniscus of fluid that forms in imaging chambers. As shown in Figure 2A, deconvolution of phase density (Phase) from brightfield (BF) data (11, 35) and deconvolution of fluorescence density (FL density) from raw fluorescence (FL) improves the contrast for organelles. The visualization of organelles is improved because spatial frequencies that represent organelle texture are typically within the middle of the passband of optical microscopes. These spatial frequencies are transmitted to the image, but also suppressed. The deconvolution corrects this imbalance. The deconvolved phase density also reports the local dry mass of the cells more consistently. In brightfield images, dense structures are transparent in focus, and brighter or darker relative to the background when out of focus. In the deconvolved phase density images, the contrast is more uniform (Figure 2A). The model trained to predict fluorescence density from phase density leads to the sharpest predictions of nuclei and membrane. This is reflected in higher average precision (AP) and higher AP at the IoU (Intersection over Union) threshold of 0.5 (AP@0.5) between the instance segmentations of nuclei from fluorescence density images and from virtually stained images (<u>Figure S3</u>).

**Figure 2.**
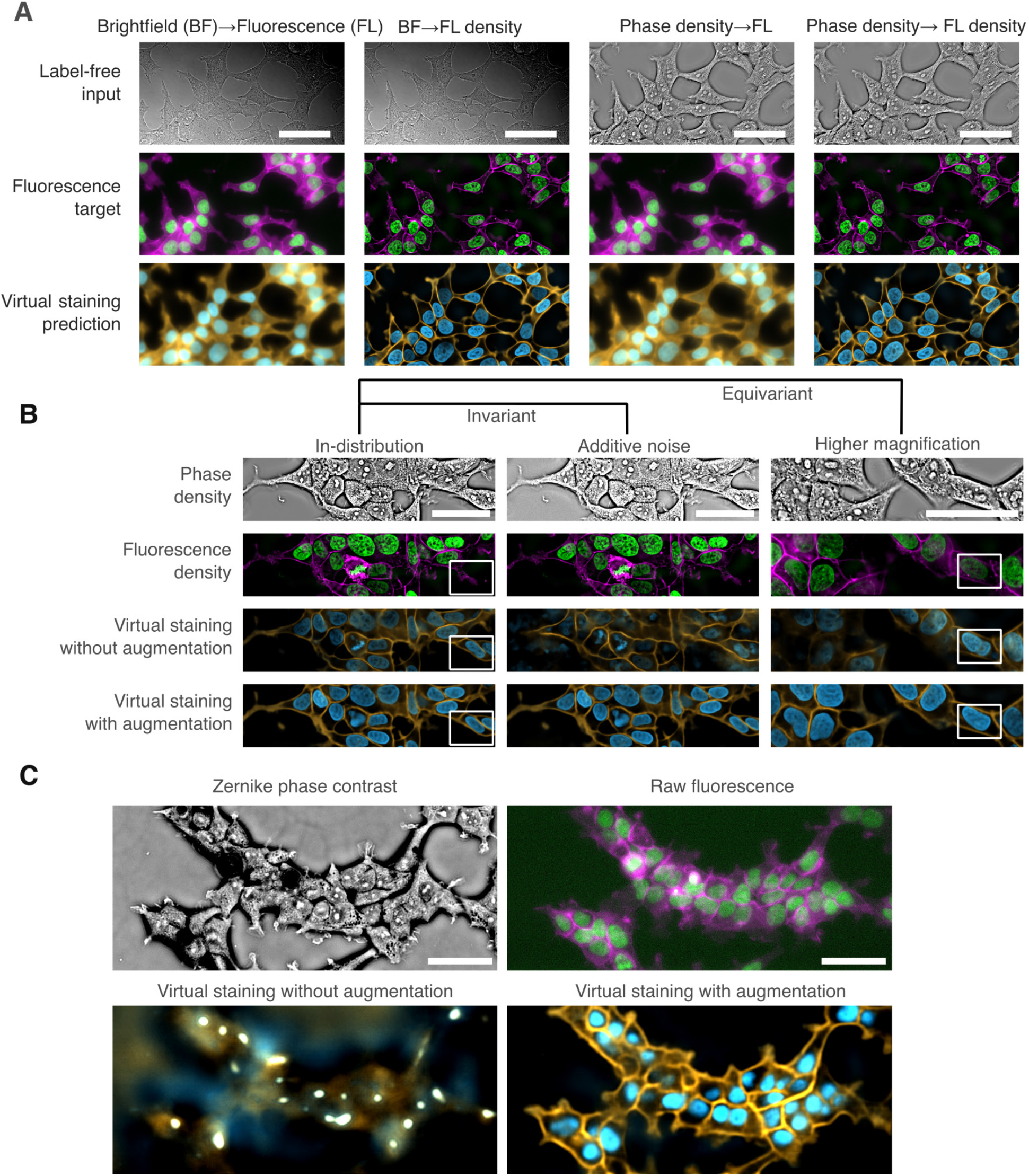
Deconvolution and data augmentation strategies make the VSCyto3D model robust to label-free imaging parameters: **(A)** Physics-based reconstructions enhance contrast for virtual staining. Top to bottom: label free input, fluorescence target, and virtual staining prediction. Models are trained on pairs of raw or reconstructed label free and fluorescence contrast modes. Scale bars: 50 µm. **(B)** Predictions of nuclei and membrane from phase image (1st row) using models trained without augmentations (3rd row) are inconsistent with experimental ground truth (2nd row), especially in the presence of noise (center column) or at a different magnification (right column). The predictions using the models trained using spatial and intensity augmentations (see text for details) are invariant to noise and equivariant with magnification. The white box in the in-distribution column highlights the rescue of the lost fluorescence label. The white box in the higher magnification column shows that the model with augmentations correctly predicts one large nucleus whereas the model trained without augmentation predicts two smaller nuclei. Scale bars: 50 µm. **(C)** Data augmentation improves generalization to unseen modality. Virtual staining models were trained to predict fluorescence density from phase density and then used to predict nuclei and plasma membrane from Zernike phase contrast image (top left). The correlative raw fluorescence image (top right) shows low signal-to-noise ratio due to light loss in the phase contrast objective. Scale bars: 50 µm.

Interestingly, the deconvolution causes a drop in Pearson correlation coefficient (PCC) between virtually stained and fluorescence density images, because the virtually stained fluorescence is inherently smooth (<u>Figure S3</u>). The smoothing enables robust segmentation of landmark organelles in the presence of noise, but is detrimental for virtual staining of structures close to the resolution limit of the microscope. The improvement in segmentation metric, but worsening of regression metric due to deconvolution of data illustrates the need for nuanced interpretation of the metrics.

Data augmentations that account for the formation of natural and medical images have been important for robust representation learning (43) and segmentation (44). We reasoned that training data should be augmented with spatial and intensity filters inspired by the image formation of microscopes to make the predictions of our models invariant to exposure, noise, the size of the illumination aperture, and similar nuisance variations in imaging parameters. Figure 2B illustrates the images without and with such spatial and intensity augmentations (<u>Methods-Data Augmentations</u>). The predictions (Figure 2B, virtual staining with augmentations) and segmentations (<u>Figure S4</u>) across the test dataset become invariant to variations in imaging parameters as we incorporate spatial and intensity augmentations inspired by image formation. As expected, the scaling augmentations make the model equivariant to magnification.

Fluorescent labeling is stochastic, especially when cells are engineered to express multiple fluorescent tags (7). Sampling the patches from the training data in proportion to the degree of labeling makes the models robust to partial and uneven labeling as shown in Figure 2B (white box). In fact, the VSCyto3D model rescued the nuclear stain in the fields of view from the test dataset (<u>Figure S5</u>) where many cells were missing the nuclear stain. Comparison of the 3D distribution of experimentally and virtually stained nuclei and membrane in a through focus movie (<u>Video 1</u>) shows that virtual staining improves the uniformity of labeling of cell membrane.

We also explore if the label-free input images can be simulated to mimic acquisition with a different light path that transfers less information than the light path with which the training data was acquired. Filters informed by the image formation process were included in the augmentation pipeline for input data to simulate the data acquisition with Zernike phase contrast (PhC). This strategy enabled generalization of the VSCyto3D model to PhC images (Figure 2C) not seen during the training. The raw fluorescence images of labeled nuclei and membranes acquired with the PhC objective were significantly blurrier and noisier (Figure 2C, raw fluorescence) than those acquired with the widefield objective, due to the phase ring specific to the PhC objective that absorbs and filters fluorescence emission. Interestingly, virtually stained nuclei and membranes are sharper (Figure 2C, virtual staining with augmentation), and therefore lead to sharper instance segmentations of nuclei and membranes (<u>Figure S6</u>). In other words, the augmentations we devised expanded the input image space to include Zernike phase contrast-like images, while constraining the space of possible output images to sharp fluorescence density images. This strategy also enabled synthesis of training datasets at 20x magnification for training VSCyto2D model (Figure 1B, <u>Methods-Model</u> <u>Training</u>).

The degree of perturbation to which the model is robust was assessed by simulating the blur and contrast stretch in the input image. The VSCyto3D model performs well, even when the images show significant blur and contrast variation (<u>Figure S7</u>). This implies that the model is robust to variations in numerical aperture that modulate the resolution and contrast of a phase image. In order to spot-check that the VSCyto3D model learns a meaningful mapping between imaging modalities, we visualize the feature maps (<u>Methods</u> - <u>Model visualization</u>) at each level of encoder and decoder (<u>Figure S8</u>) for a randomly chosen test image. The boundaries of cells and nuclei can be identified at higher levels of abstraction in the encoder and the decoder.

These results demonstrate a training protocol for robust virtual staining that consists of acquiring training data at the highest possible resolution, deconvolving it with an image formation model, augmenting with filters to mimic the changes in contrast and resolution, and sampling it in proportion to the degree of labeling.

### Virtual staining of evolving 3D organs

Another key challenge in virtual staining is to accurately predict landmark organelles during morphological changes associated with development and differentiation. One such example are the neuromasts of the zebrafish lateral line, whose three-dimensional shapes and textures change throughout their development (9, 10). Using zebrafish neuromasts as a model system, we explore an economical strategy to generalize 3D virtual staining models (VSNeuromast) to different developmental stages. Moreover, the training protocol provides robustness to imaging challenges characteristic of in-vivo time-lapse experiments.

We trained a baseline VSNeuromast model with UNeXt2 architecture with 21 z-slices as input and output (Figure 1, <u>Figure S9</u>) following the training protocol described for the VSCyto3D model, using data acquired on the widefield fluorescence microscope at two developmental stages (3dpf and 6.5dpf, dpf = days post fertilization). When this model was used to predict nuclei and membrane at 4 dpf and compared with confocal images of nuclei and membrane, no hallucinations were noticed, but the predictions were blurry (Figure 3A, row: virtual staining without fine-tuning). Images at 4dpf are slightly out of the distribution of the training data acquired at 3dpf and 6.5dpf.

**Figure 3.**
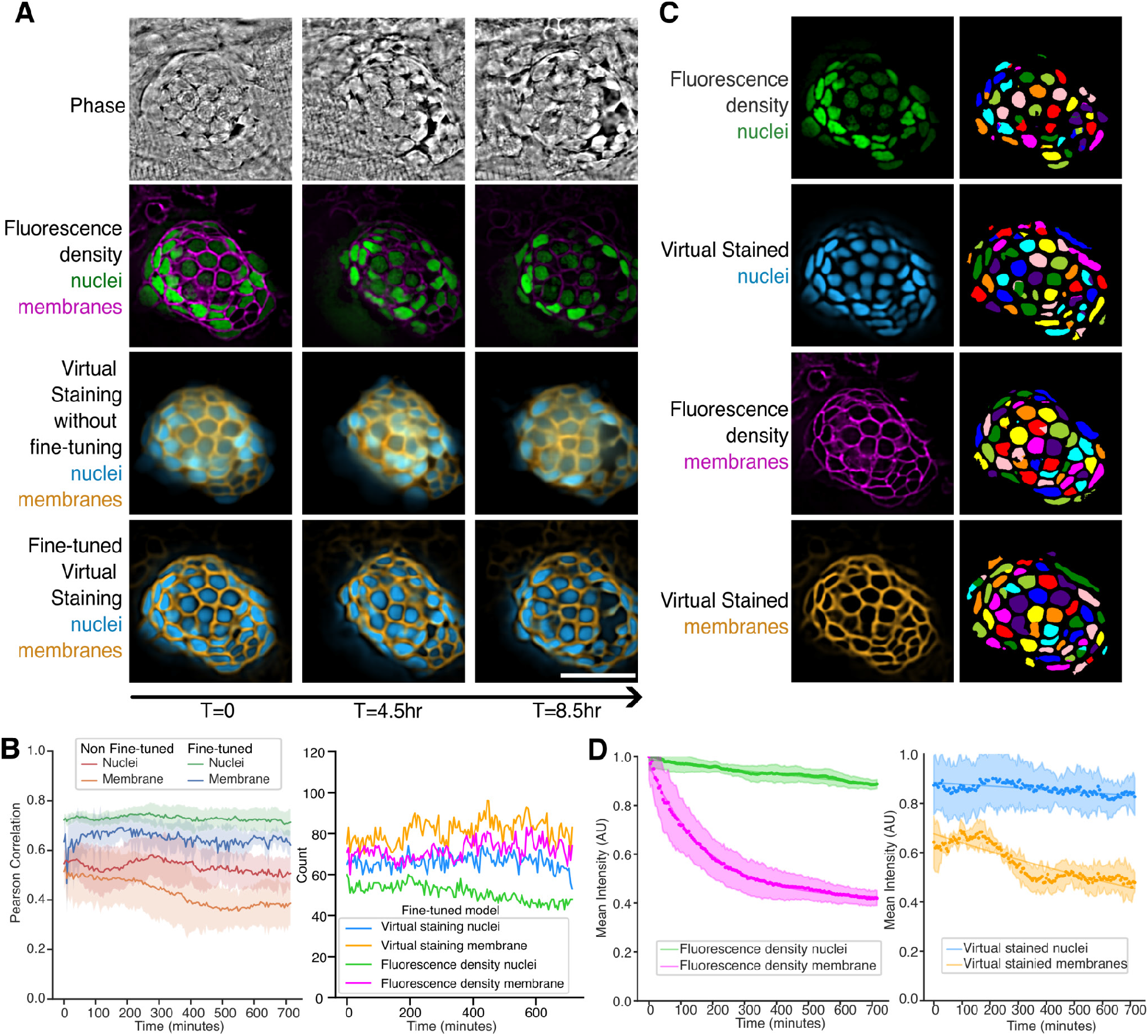
Generalizing the VSNeuromast model across zebrafish development **(A)** Phase (1st row), experimentally stained nuclei and membrane (2nd row), and virtually stained nuclei and membrane using a model that was not fine-tuned (3rd row) and using a model that was fine-tuned on the subsampled movie (4th row) are shown. We show 3 samples from a 12-hour movie starting at 4 days post-fertilization (4 dpf) and microscope. The VSNeuromast model was fine-tuned with subsampled 4dpf movie (1/12 timepoints). Virtual staining rescues missing nuclei and provides a more accurate read-out of the cell count and their locations than experimental staining. (Scale bar 25um) **(B)** Comparison of nuclear and membrane 3D segmentations predicted using the fine-tuned VSNeuromast Cellpose model. The same segmentation model was applied consistently to fluorescence density and virtual staining volumes. The VSNeuromast and Cellpose segmentation combination predicts the cell counts with high accuracy. The model’s performance was quantitatively assessed using Pearson correlation plots across five neuromasts from the lateral line, comparing both fluorescence density and virtual staining results for nuclei and membranes to highlight the precision of the fine-tuned model. **(C)** Segmentation of nuclei and membranes using fine-tuned Cellpose model on experimental fluorescence and virtual staining. Virtual staining reduces over- and under-segmentations, enhancing accuracy compared to experimental fluorescence. **(D)** Mean photobleaching curves across five neuromasts showing experimental fluorescence (left) and virtual staining (right) nuclei and membrane pairs. The shaded region indicates the variation in the mean intensity of nuclei in a given frame.

An economical strategy for generalization of virtual staining across developmental time without the risk of hallucinations is to fine-tune a pre-trained model with coarsely sampled pairs of label-free and fluorescence data. With this strategy, nuclei and cells can be tracked at high temporal resolution with phase imaging and the model’s accuracy can be continuously calibrated. This approach can reduce the photodamage, enabling faster and longer imaging of developmental dynamics. To assess the viability of this approach, the movie of neuromasts imaged at 4dpf is subsampled by the factor of 1/12 to create a fine-tuning training set. The fine-tuning workflow is described in <u>Methods-Model Training</u> and <u>Figure S10</u>. Visualization of predictions on a neuromast from the validation set (<u>Video 2</u>) illustrates how blurry predictions of the pre-trained model are sharpened during fine-tuning.

The fine-tuned model is more accurate (Figure 3A, row: fine-tuned virtual staining). As shown in Figure 3B, fine-tuning improves the PCC between the virtually stained nuclei and membrane and experimentally stained nuclei and membrane. The fine-tuned model enables robust virtual staining of nuclei and membrane in 3D and over time as seen from the comparison of experimental and virtually stained neuromast in <u>Video 3</u>.

The virtually stained nuclei show a more uniform intensity distribution than the experimentally stained nuclei as seen from Figure 3A and Figure 3C. The dimmer nuclei in the fluorescence density image are missed by the segmentation model but are rescued by virtual staining (<u>Video 4</u>). A comparison of the mean cell count over time obtained from fluorescently labeled and virtually stained nuclei of five lateral line neuromasts corroborates the rescue (Figure 3B, cell count, green curve vs blue curve) of dim nuclei seen in <u>Video 4</u> and Figure 3C. A comparison of instance segmentation from experimentally and virtually stained membrane shows less pronounced, yet measurable, rescue of cells by virtual staining (Figure 3B, cell count, magenta curve vs orange curve). In both experimental and virtually stained neuromasts, we notice extraneous cell segmentations at the edge of neuromasts. These extraneous segmentations can be filtered via tracking.

The mean intensity of each segmented cell’s nuclei and membrane were compared (Figure 3D) to assess whether the model can rescue photobleaching. The experimental membrane channel shows clear photobleaching. On the other hand, the photobleaching is measurably, but not fully, corrected by the fine-tuned virtual staining model. After further refinement, we anticipate that virtual stain can substitute the experimental stain and reduce photobleaching/photodamage.

Interestingly, the VSNeuromast model virtually stains cells around the yolk, because the size and texture of these cells resemble cells of neuromast. These cells can be easily filtered in post-processing as shown in <u>Figure S11</u>. This data, however, suggests a possibility of training a model that virtually stains all nuclei in zebrafish if the phase images are acquired with sufficient resolution.

The features learned by the VSNeuromast model were interpreted by visualizing the feature maps learned by the encoder and decoder (<u>Methods-Model Visualization</u>). The model indeed represents shapes of nuclei, membrane, and neuromast as seen from the principal components of the feature maps shown in <u>Figure S12</u> for an example input image of neuromast. These results show that the fine-tuning of a virtual staining model on subsampled pairs of phase and fluorescence volumes enables generalization across developmental stages and can mitigate the photodamage caused by high-temporal fluorescence imaging.

### Few-shot generalization to new cell types

Next, we report generalization of robust virtual staining models to new cell types with minimal new training data using a pre-training/fine-tuning paradigm common in language and vision modeling. Collecting large amounts of paired label-free and fluorescence images with sufficient diversity of cell morphology is often difficult. For example, consistent labeling of cell membranes requires genetically expressed peptides (e.g., CAAX) that localize to cell membranes. Engineering cells to express genetic labels is time-consuming and challenging in cells that are not immortalized. Since the landmark organelles exhibit common morphological features across cell types, we reasoned that adapting the pre-training/fine-tuning protocol developed for image classification (40) to image translation can enable few-shot generalization of virtual staining models to a new cell type.

We explore generalization of the models for virtual staining of nuclei and cell membranes in HEK293T and A549 to two new cell types: BJ-5ta, immortalized fibroblast cells used in toxicology research, and iNeurons derived from KOLF2.1J-NGN2 human induced pluripotent stem cells (iPSCs), used in neurobiology research. Virtual staining of BJ-5ta cells will be used for image-based screening of cellular impacts of viral infection. Virtual staining of iNeurons will be used for label-free quality control (QC) of the differentiation process. Maintenance and differentiation of iPSCs takes weeks and requires diligent care. Due to high batch-to-batch variability, robust QC is essential to ensure reproducible differentiations and measurements. QC of iNeurons involves evaluating the morphology of the cells to ensure they exhibit the expected neuronal phenotype, including the presence of cell bodies, neurites, and axons. The neuronal phenotype is typically evaluated with the following morphological features: 1) cell bodies exhibit a characteristic round or polygonal shape with prototypical size and a centrally located nucleus; 2) Mature neurons exhibit neurites, including axons and dendrites, projecting from the cell body.

The computational experiments described next use 2D images at lower magnification (see Method-Model Training for details) common in image-based screens and result in the VSCyto2D model.

Figure 4A illustrates a self-supervised pre-training and supervised fine-tuning protocol, which uses images of HEK293T and A549 cells for pre-training virtual staining models that are fine-tuned for virtual staining of BJ-5ta and iNeuron cells. During self-supervised pre-training (Figure 4B), the phase images are randomly masked, and the unmasked pixels are used to predict the masked pixels in each training patch (<u>Methods-Model Training</u>), following the fully convolutional masked autoencoder (FCMAE) protocol reported for image classification (40). The model is pre-trained in two steps: 1) The encoder and decoder weights are optimized with just phase images of HEK293T and A549 cells using the masked autoencoding task shown in Figure 4B, 2) The weights are transferred to a virtual staining model that is pre-trained to predict fluorescent nuclei and cell membranes using HEK293T and A549 cells. <u>Video 5</u> shows that the model pre-trained with HEK293T and A549 datasets generalizes well to diverse cell shapes of A549 cells observed throughout the cell cycle. After the pre-training, the model is fine-tuned with data acquired with a new cell type (BJ-5ta or iNeuron) that has a distinct morphology. The computational graphs of the models used for pre-training and fine-tuning are shown in <u>Figure S13</u>.

**Figure 4.**
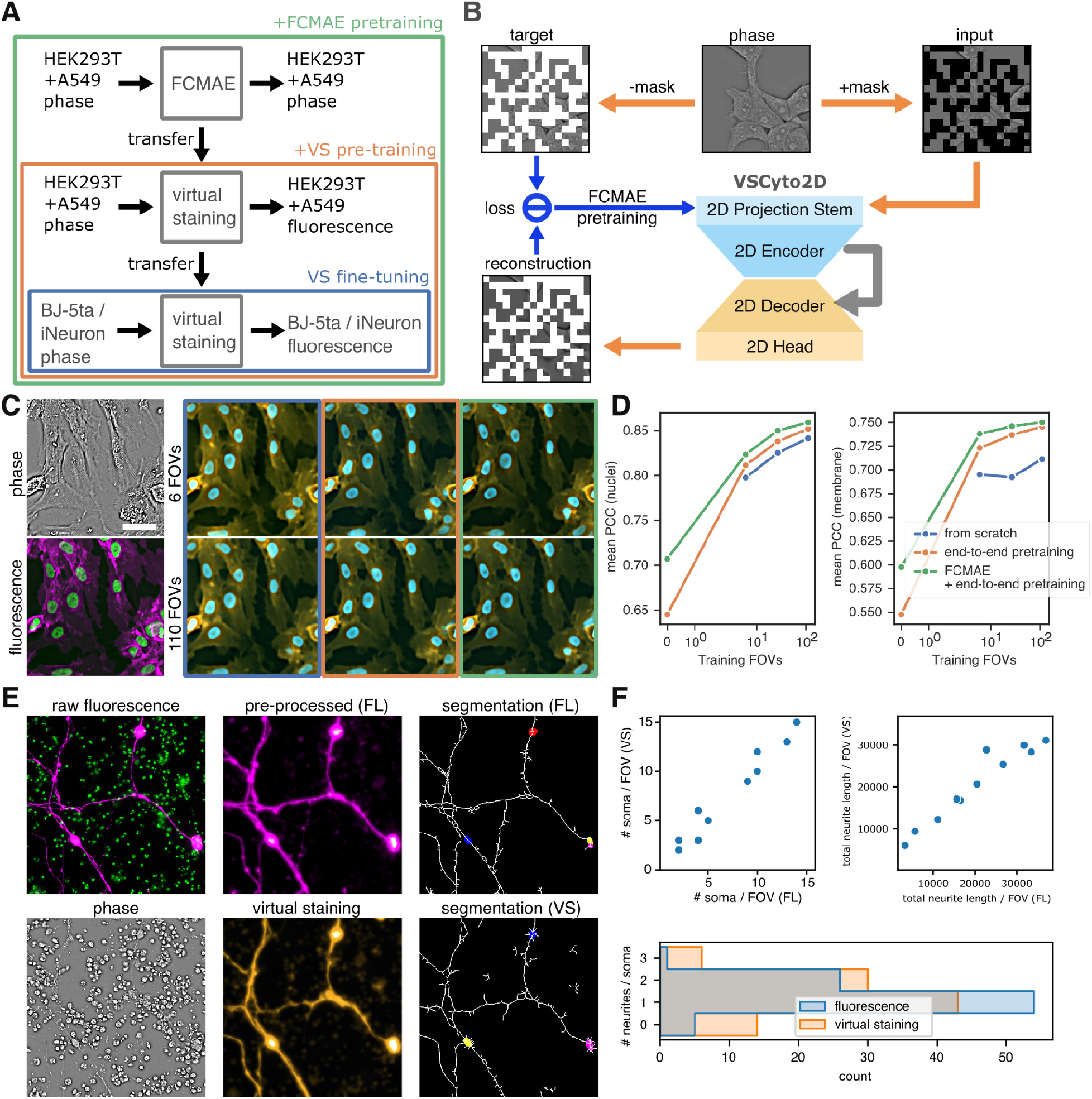
Few-shot generalization of the VSCyto2D model to new cell types **(A)** The encoder of UNeXt2 is pre-trained with a fully convolutional masked autoencoder (FCMAE) recipe to enable generalized feature encoding without paired data. The model can then be fine-tuned for virtual staining on a specific cell type. **(B)** Flow chart of 3 training strategies to generalize the virtual staining model pre-trained with A549 and HEK293T cells to the BJ-5ta cell type with limited training samples. The bounding boxes indicate strategies: (blue) virtual staining pre-training from scratch with paired images of BJ-5ta; (orange) pre-training with paired images of HEK293T and A549, and fine-tuning with paired images of BJ-5ta; and (green) FCMAE pre-training with only the phase images of HEK293T and A549, virtual staining pre-training with images of HEK293T and A549, and fine-tuning with paired images of BJ-5ta. The pre-training steps initialize model weights in the encoder (FCMAE) and decoder (virtual staining) of UNeXt2. **(C)** Virtual staining images of nuclei and membrane in BJ-5ta using 3 models described in D. Performance scales with the increasing number of BJ-5ta FOVs used for fine-tuning. Scale bar: 50 µm. **(D)** Pearson correlation coefficient (PCC) between experimental and virtually-stained nuclei and plasma membrane as a function of the number of fields of view used for the test dataset used for training strategies shown in B. The pre-trained models show superior performance scaling relative to the number of BJ-5ta FOVs used for fine-tuning. Pre-trained models fine-tuned with less data can match or outperform models trained with more data from scratch. **(E)** Virtual staining of iNeuron cells recovers contrast needed for soma segmentation and neurite tracing. Soma segmentation is shown in color-filled labels, neurite tracing is shown in white lines. Scale bars: 100 µm. **(F)** Quantitative analysis of iNeuron segmentations. Similar number of soma per FOV, total neurite length, and number of neurites per soma can be identified from virtual staining compared to experimental staining.

When using data from just one cell type (HEK293T), the pre-training/fine-tuning protocol slightly improves the visual sharpness of predicted images as shown in <u>Figure S14</u>. As shown earlier (<u>Figure S5</u>), some of the fields of view had missing nuclei labels. These fields of view were proofread, and segmentation and regression metrics (<u>Figure S14</u>) were computed from virtually stained and experimental fluorescence images. These metrics show that the models pre-trained on label-free images with masked autoencoder, and fine-tuned on paired data are as accurate as models trained from scratch.

Figure 4C reports few-shot generalization to BJ5-ta cells. The images (Figure 4C) and segmentations (<u>Figure S15</u>) show that the pre-trained model after being fine-tuned with 6 fields-of-views (FOVs) performs as well as the model trained from scratch on with ∼100 FOVs. Visualization of the evolution of the predictions from the validation set (<u>Video 6</u>) for the models trained with different training protocols show that pre-trained models produce useful predictions from the first epoch. Comparing the segmentation metrics for nuclei and membrane as a function of the number of training FOVs (Figure 4D) confirms that pre-trained/fine-tuned models scale better, i.e., generate more accurate predictions, relative to the models trained from scratch.

The effect of different training protocols on features learned by the models is illustrated by visualizing the feature maps of the models (<u>Methods-Model Visualization</u>). We find that the model pre-trained on phase images (<u>Figure S16</u>, column 3, rows: encoder stages) learns a more regular representation of cell boundaries than the models trained on just the virtual staining task (<u>Figure S16</u>, columns 1 and 2, rows: encoder stages).

Figure 4E reports fine-tuning of the VSCyto2D model to predict the soma and neurites of iNeurons from phase images. The images acquired with vital dyes (Figure 4E, raw fluorescence) that stain nuclei and live cells were pre-processed (<u>Methods-VSCyto2D</u>) to suppress the dead cells. In this case, the cells that didn’t attach to the substrate at the start of differentiation died. The pre-processing step synthesizes clearer contrast for neurites (Figure 4E, pre-processed, magenta) and for soma (Figure 4E, pre-processed, green). The synthesized fluorescence channels were used as a target for fine-tuning the VSCyto2D model. The fine-tuned model enables detection of soma and neurites (Figure 4E, virtual staining) even in the presence of dead cells in the field of view (Figure 4E, phase). The utility of the virtual staining model for QC of differentiation is assessed by segmenting the soma and neurite from pre-processed fluorescence images or virtually stained images, and computing the following metrics of neuronal phenotype (Figure 4F): number of live soma per field of view, total length of neurites within a field of view, and the number of neurites/soma. The features retrieved from virtually stained images corroborate the features retrieved from pre-processed fluorescence images. The model achieved this robustness with a training and validation set consisting of ∼500 iNeurons, in contrast to ∼11000 HEK293T and A549 cells used during pre-training. Taken together, the above results establish a training protocol for generalizing virtual staining models to new cell types.

Above results demonstrate several new strategies for virtual staining of cellular landmarks across imaging conditions, cell states, and cell types. The physics-informed data augmentations enabled zero-shot generalization of the VSCyto3D model to Zernike phase contrast without the need to acquire training data with this modality. These augmentations also made the model robust to nuisance factors such as non-uniform illumination and changes in numerical aperture. This robustness is particularly critical for practical image-based screens that integrate data from diverse microscopes with varying imaging conditions and optical aberrations. The fine-tuning of the model using subsampled time series led to generalization of the VSNeuromast model to different developmental stages. Lastly, the pre-training/fine-tuning protocol enabled few shot generalization of the VSCyto2D model to new cell types. This strategy leverages the consistency of organelle shapes across different cell types, significantly reducing the data requirements for training robust virtual staining models. The performance improvement achieved by pre-training with a masked autoencoding task followed by fine-tuning suggests a viable strategy for scaling virtual staining models to a broader range of biological samples.

We report a diverse set of evaluation metrics, including regression metrics, instance segmentation metrics, and application-specific measurements to evaluate the models’ performance for real-world biological research. Inspection of learned features confirms that the data augmentation strategies and training protocols described guide the model in learning a semantic mapping of cell structures between input and target imaging modalities. This capability is fundamental for the accurate virtual staining of cellular structures. We also illustrate failure modes and regime of validity of each of the three models reported here.

This work paves the way for the following developments in virtual staining and its applications. First, simulations with image formation models may further generalize the models to other phase imaging modalities without the need to acquire new data. Second, the test-time augmentations may make predictions of our models even more robust. Third, the pre-training/fine-tuning strategy may be extended to train decoders for landmark organelles other than nuclei and cell membranes, such as nucleoli and lipid droplets. Fourth, the pre-training strategy can be used across developmental stages of zebrafish, enabling label-free tracking of cells across developing embryos. Finally, the training protocols developed for virtual staining can be adapted for segmentation models, potentially leading to joint virtual staining and segmentation models that offer even greater generalizability and accuracy.

We share the above models and the pipeline for training, evaluating, and deploying the models via our GitHub repository, VisCy and anticipate that our work will accelerate the mapping and understanding of dynamic cell systems.

### Image acquisition and dataset curation

#### Human cell lines (HEK293T, A549, BJ-5ta, and iNeuron)

HEK293T cells are labeled with H2B-mIFP and CAAX-mScarlet following the OpenCell protocol (45). Brightfield and fluorescence volumes of live HEK293T cells cultured in a 24-well plate are acquired on a wide-field fluorescence microscope (Leica Dmi8) with a 63x magnification, 1.3 NA glycerol-immersion objective to collect training data. For testing robustness to different imaging conditions shown in Figure 2, additional volumes are imaged with a 40x magnification, 1.1 NA water-immersion objective, and a 100x magnification, 1.47 NA oil-immersion objective, at 0.25 µm Z-steps, for 96 Z-slices. For testing generalization to the Zernike phase contrast modality, image volumes are acquired with a 40x magnification, 0.6 NA Ph2 air objective, at 0.4 µm Z-steps, for 58 Z-slices. The images are sampled on a camera with 6.5 µm pixel size and 2048x2048 sensor For training, the Z-slice corresponding to the coverslip is determined by maximizing the transverse mid-band power of the mScarlet fluorescence density channel, and the Z-slices ranging from -2 µm to +12.5 µm relative to the coverslip were kept. Volumes that do not contain this range are excluded from the training dataset. The training dataset contains 291 volumes. For testing robustness to imaging conditions, images were acquired on a different day with new cell cultures, and 12 volumes were acquired for each condition.

A549 and BJ-5ta cells were cultured in 12-well plates and stained with Hoechst for nuclei and CellMask for plasma membrane. The A549 cells were cultured in Dulbecco’s modified eagle medium (DMEM). The cells were seeded at 1.3⨉10^5^ cells/well in a 12 well glass bottom plate and imaged after 31 hours. DMEM was replaced with imaging media right before imaging.

Human KOLF2.1J-NGN2 iPSC cells were differentiated into iNeurons following (46), seeded in a 24-well plate in differentiation media, and imaged at 7 days post differentiation. To determine whether our virtual staining pipeline can be used for QC of iNeurons, we stained iNeurons with Hoechst and Calcein, a non-invasive fluorescent dye, to assess cell viability and neuronal health. Calcein was chosen because it specifically labels live cells, allowing for easy monitoring of viability across the neuronal culture. Labeling depends on enzymatic activity, serving as an indicator of cell health and metabolic function, critical for efficient differentiation. Furthermore, Calcein reports cell morphology, which is commonly used to assess neuronal differentiations. Healthy neurons should exhibit morphological features e.g. neurite outgrowth and network formation, which are easily visualized by Calcein. In our hands, Calcein started to affect the health of iNeurons within 6 hours of staining. We used only healthy cells for training the virtual staining models.

Brightfield and fluorescence volumes of live cells were acquired on a wide-field fluorescence microscope (Leica Dmi8) with a 20x magnification, 0.55 NA objective. The images were sampled on a camera with 6.5 µm pixel size and 2048 by 2048 pixel sensor size, at 1 µm Z-steps. Each dataset was split into a training and validation subset and a testing subset as shown in the following table:

| Cell type | Training and validation FOVs | Testing FOVs |
| --- | --- | --- |
| A549 | 24 | 7 |
| BJ-5ta | 138 | 12 |
| iNeuron | 70 | 11 |

An additional A549 dataset (<u>Video 5</u>) was acquired to test the generalization of the VSCyto2D model to dynamic cell states. Time-lapse imaging was performed at a time interval of 1 hour, for a total of 17 hours.

#### Zebrafish neuromasts

Following the approved IACUC protocols, this study utilizes transgenic zebrafish lines expressing she:H2B-EGFP and cldnb:lyn-mScarlet (47) to label the nucleus and cell membrane, respectively, of the neuromasts. The VSNeuromast model utilizes neuromasts collected from three developmental stages: 3, 6 and 6.5 dpf (days post-fertilization: dpf). The datasets are composed primarily of lateral line neuromasts resulting in a dataset with 273 total volumes (160 volumes from 3dpf, 57 from 6dpf and 56 from 6.5dpf) with ZYX shape (107, 2048, 2048) or (35.3µm, 237.6µm, 237.6µm). The dataset was split 80/20 for training and validation respectively. Additionally, 6 neuromasts from an acquisition from a different fish and day are used as a test dataset. For each neuromast, a brightfield and two fluorescence channel stacks were acquired at Nikon PlanApo VC x63 1.2NA objective on an ASI Rapid Automated Modular Microscope System (RAMM) through the same optical path using a Andor ZYLA-4.2P-USB3-W-2V4 sCMOS camera. Channels were well registered because they shared the same imaging path. The fluorescence stacks of labeled nuclei and cell membrane were used as target channels.

The VSNeuromast model is fine-tuned for Figure 3 using the same transgenic line. Five lateral line neuromasts are imaged with an Olympus IX83 dual turret microscope system using a 63x objective for fluorescence and label-free imaging every 5 minutes over a total of 12hrs. The VT-iSIM system with a Hamamatsu Quest v1 C15550-20UP qCMOS camera with 4.6 µm pixel size is used for the fluorescence imaging. The label-free imaging uses a custom imaging path built on the first level of the microscope body using a 200mm tube lens (Thorlabs TTL-200A MP) resulting in 66x effective magnification using a machine vision camera (BFS-U3-51S5M-C). The test dataset was acquired with a temporal interval of 5 minutes and the fine-tuning dataset is created by subsampling the timelapse by using the volumes acquired every hour. The fine-tuning datasets are split 80/20 into training and validation datasets.

#### Data conversion

All internal datasets are acquired in uncompressed lossless formats (i,e OME-TIFF and ND-TIFF) and converted to OME-Zarr (48) using iohub (49), a unified python library to convert from common bio-formats (i.e OME-TIFF, ND-TIFF, Micro-Manager TIFF sequences) and custom data formats to OME-Zarr.

### Preprocessing

#### Deconvolution of phase density and fluorescence density

The reconstruction from brightfield and fluorescence stacks to phase density and fluorescence density is performed with the <u>recOrder</u> library (11, 35, 50). Briefly, the acquired brightfield and fluorescence stacks are modeled as filtered versions of the unknown specimen properties, phase density and fluorescence density, respectively. This blur is represented by a low pass optical transfer function (OTF) in Fourier space and a point spread function in the real space, which are simulated using properly calibrated parameters of the imaging system (numerical apertures of imaging and illumination, wavelength of illumination, and pixel size at the specimen plane). The simulated OTFs are used to restore phase density and fluorescence density, respectively, from the brightfield and fluorescence stakes using a Tikhonov-regularized inverse filter.

#### Registration

The label-free and fluorescence channels are registered with shrimPy (51). The resulting volumes are cropped to ZYX shape of (50, 2044, 2005) for the HEK293T Zernike phase contrast test dataset, (9, 2048, 2048) for A549, (12, 2048, 2009) for BJ-5ta, and (26, 2048, 2007) for iNeuron. The neuromast datasets acquired with the wide-field fluorescence microscope are registered to the phase density channel and cropped to (107, 1024, 1024). The datasets acquired in the iSIM setup are cropped to (81, 1024, 1024).

#### Additional preprocessing (iNeuron)

The fluorescence signal in iNeuron cells was further processed to improve contrast for virtual staining and segmentation.

For the Calcein channel, the soma is much brighter than the neurites. Mean projection along the axial dimension and natural logarithm of one plus the input (‘log1p’) are applied to compress the dynamic range. The result is normalized so that the 99th percentile is 0, and the 99.99th percentile is 1, and then clipped to a range of 0 to 5.

To suppress fluorescence from dead cells in the Hoechst channel, the max projection of Hoechst volumes is multiplied with the mean projection of the raw Calcein channel. The result is normalized so that the median is 0, and the 99.99th percentile is 1, and then clipped to a range of 0 to 5.

To match the shape of the fluorescence channels, a single Z-slice (at 8 µm from the bottom of the volumes) is taken from the phase channel as the input to virtual staining models.

### Model architecture

We use an asymmetric U-Net model with ConvNext v2 (40) blocks for both virtual staining and FCMAE pre-training. The original ConvNext v2 explored an asymmetric U-Net configuration for FCMAE pre-training and showed that it has identical fine-tuning performance on an image classification task. In the meantime, SparK (41) used ConvNext v1 blocks in the encoder and plain U-Net blocks in the decoder for its masked image modeling pre-training task. We use the ‘Tiny’ ConvNext v2 backbone in the encoder. For FCMAE pre-training, 1 ConvNext v2 block is employed per decoder stage. For virtual staining models, each decoder stage consists of 2 ConvNext v2 blocks.

### Model training

Intensity statistics, including the mean, standard deviation, and median were calculated at the resolution of FOVs and at the resolution of the whole dataset by subsampling each FOV using square grid spacings of 32 pixels in each camera frame. These pre-computed metrics are then used to apply normalization transforms by subtracting the choice of median or mean and dividing by the interquartile range or standard deviation respectively. This enables z-scoring of the training data at the level of the whole dataset, at the level of each FOV, and at the level of each patch (11), depending on the use case.

#### Training objectives

The mixed image reconstruction loss (52) is adapted as the training objective of virtual staining models: 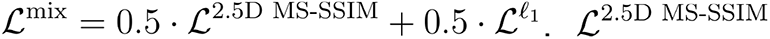 is the multi-scale structural similarity index (53) measured without downsampling along the depth dimension, and 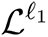 is the L1 distance (mean absolute error). The virtual staining performance of different loss functions is compared in Table 2.

The mean square error (MSE) loss is used to pre-train the FCMAE models on label-free images, following the original FCMAE implementation (40).

#### Data Augmentations

The data augmentations are performed with transformations from MONAI (44).

##### Spatial augmentation

| MONAI transformation | Parameters |
| --- | --- |
| Random scaling | $\pm 0.3$ or $\pm 0.5$ in XY, and $\pm 0.2$ in Z. |
| Random rotation | $\pm \pi$ around the Z axis. |
| Random shearing | $\pm 0.05$ in XY |
| Random XY flip | only used for VSCyto2D |

##### Intensity augmentations

| MONAI transformation | Parameters |
| --- | --- |
| Random contrast adjustment | gamma range (0.8, 1.2) |
| Random intensity scaling | $\pm 0.5$ |
| Random Gaussian blur: | sigma range (0.25, 0.75) in XY |
| Random Gaussian additive noise | strength determined per dataset (sigma 0.3-5). |

#### VSCyto3D

##### Normalization

For each channel in the HEK293T dataset, the image volume is subtracted by its dataset level median and divided by the dataset level interquartile range.

##### Training

Models are trained with a warmup-cosine-annealing schedule on 4 GPUs with the distributed data parallel (DDP) strategy. A mini-batch size of 32 and learning rate of 0.0002 is used. Training and validation patch ZYX size is (5, 384, 384). For testing the effect of deconvolution (Figure 2B), models are trained for 100 epochs. For testing robustness to imaging conditions (Figure 2D), models are trained for 50 epochs.

#### VSNeuromast

##### Data pooling

The data used in our methods is pooled from four OME-Zarr stores, which contain neuromasts from 3dpf, 6dpf, and 6.5dpf stages. These stores include both the whole field of view (FOV) and a center-cropped version focused on the neuromast. For the cropped FOVs, a weighted cropping technique is applied to ensure the inclusion of training patches containing the neuromast. Conversely, the uncropped dataset employs an unweighted cropping method to incorporate additional contextual information. A high content screening (HCS) dataloader was developed to sample equally from the multiple datasets with variable length.

In the fine-tuning step, the experimental fluorescence channels were registered to the phase density channel and required downsampling of the data by the factor of 2.1 to match the pixel size between the datasets used for pre-training and fine-tuning.

##### Normalization

This model normalizes the label-free channel per FOV by subtracting the median and interquartile range.

##### Training

The models are trained with a warmup-cosine-annealing schedule on 4 GPUs with the distributed data-parallel (DDP) strategy. This model uses datasets from 3dpf and 6-6.5dpf using mini-batch size of 6 from each dataset and learning rate of 0.002.

The VSNeuromast model is initially trained using ZYX patch size of (15,384,384) for 150 epoch. The weights from this model are loaded to train a model that takes ZYX patch size of (21,384,384) to improve the Z prediction accuracy. Training and validation patch ZYX size is (21, 384, 384). This model is trained for 30 epochs.

##### Fine-tuning

The expression of nuclei and cell membrane labels in neuromast was equalized using a contrast adaptive histogram equalization (CLAHE) with a kernel size of [5,32,32] (z,y,x). The model is fine-tuned using the model’s checkpoint (‘neuromast_3n6dpf_21plane_v1_mixedloss_weightedcrop_hotstart_v1’) using prior patch sizes (21,384,384), learning rate 2e-4 with a warmup-cosine-annealing schedule on 4 GPUs with DDP strategy. The model is trained with 45 epochs.

The fine-tuned model is used to generate predictions shown in figures. The fine-tuned model generalizes to datasets using the same imaging setup the VSNeuromast model is trained on (Figure 1) and datasets acquired on iSIM setup described in <u>Methods</u> (Figure 3).

#### VSCyto2D

##### Data pooling

Image volumes of HEK293T cells are downsampled from the 63x dataset with ZYX average pooling ratios of (9, 3, 3). For the VSCyto2D model reported in Figure 1, training data are sampled from the downsampled HEK293T dataset, the A549 dataset, and the BJ-5ta dataset with equal weights.

##### Normalization

Each image volume is independently normalized before being used for model input. The phase channel is normalized to zero mean and unit standard deviation, and the fluorescence channels are normalized to zero median and unit interquartile range. For the iNeuron dataset, normalization is only applied for the phase channel.

##### Pre-training

FCMAE pre-training with phase images is performed with a warmup-cosine-annealing schedule for 800 epochs on 4 GPUs with the DDP strategy and automatic mixed precision (AMP). A mini-batch size of 32 and learning rate of 0.0002 was used. Mask patch size is 32, and masking ratio is 0.5. Training and validation patch ZYX size is (5, 256, 256).

##### Fine-tuning

For fine-tuning, the encoder weights are loaded from FCMAE pre-trained models when applicable. The models are then trained for the virtual staining task with the encoder weights frozen or trainable. Models are trained on 4 GPUs with the DDP strategy and AMP. Training and validation patch ZYX size is (5, 256, 256). For testing data scaling with BJ-5ta, models are trained with a constant learning rate of 0.0002. 6-FOV models are trained for 6400 epochs, 27-FOV models are trained for 1600 epochs, and 117-FOV models are trained for 400 epochs. For fine-tuning the VSCyto2D model for virtual staining of iNeurons, all parameters of the model are trained, using a warmup-cosine-annealing schedule for 1600 epochs.

##### Datasets used for model training

| Model | Reported results in | Phase data for FCMAE pre-training | Phase and fluorescence data for virtual staining pre-training | Phase and fluorescence data for fine-tuning |
| --- | --- | --- | --- | --- |
| VSCyto2D | Figure 1 | HEK293T, A549, BJ5-ta | HEK293T, A549, BJ5-ta | - |
| VSCyto2D-BJ5-ta | Figure 4 | HEK293T, A549 | HEK293T, A549 | BJ5-ta |
| VSCyto2D-iNeuron | Figure 4 | HEK293T, A549 | HEK293T, A549 | iNeuron |

##### Inference using trained models

For the 2D virtual staining model VSCyto2D, each slice is predicted separately in a sliding window fashion.

For the 3D virtual staining models (ie. VSCyto3D and VSNeuromast), a Z-sliding window of the model’s output depth and step size of 1 is used. The predictions from the overlapping windows are then average-blended.

### Model evaluation

The correspondence between fluorescence and virtually stained nuclei and plasma membrane channels are measured with regression and segmentation metrics. We describe the segmentation models for each use case below. All segmentation models are also shared with the release of our pipeline, VisCy (<u>Code and Model Availability</u>).

#### VSCyto3D

Segmentation of H2B-mIFP fluorescence density and virtually stained nuclei is performed with a fine-tuned Cellpose ‘nuclei’ model (‘CP_20220902_NuclFL’). The nuclei segmentation masks are corrected by a human annotator. Segmentation of cells from CAAX-mScarlet fluorescence density and virtually stained plasma membrane is performed with the Cellpose ‘cyto3’ model. Due to loss of CAAX-mScarlet expression in some cells, positive phase density was blended with the CAAX-mScarlet fluorescence density to generate test segmentation targets. For the Zernike phase contrast test dataset, nuclei and cells are also segmented from the phase image using the Cellpose ‘nuclei’ and ‘cyto3’ models, in addition to segmentation from experimental fluorescence images.

PCC is computed between the virtual staining prediction and fluorescence density images. AP@0.5 and mean average precision of IoU thresholds from 0.5 to 0.95 at 0.05 interval (AP) is computed between segmentation masks generated from virtual staining images and segmentation masks generated from fluorescence density images.

#### VSNeuromast

The nuclei and cell segmentations of fluorescence images are generated with fine-tuned 3D Cellpose ‘nuclei’ model and from scratch using 19 manually corrected segmentations. The segmentations are human-corrected by using the napari-annotator plugin (https://github.com/bauerdavid/napari-nD-annotator) and morphological operators such as opening, closing, and dilation to remove artifacts. The nuclei segmentation model ’cellpose_Slices_decon_nuclei_nuclei_v7_2023_06_28_16_54’ and the cell membrane segmentation model ‘’cellpose_2Chan_scratch_membrane_2024_04_01_17_12_00’ are used for neuromast segmentation across all figures.

The virtual staining models are evaluated by comparing the segmentations for fluorescence density and virtual staining predictions using Jaccard, Dice, and mean average precision (mAP) metrics at IoU thresholds of 0.5, 0.75, and 0.95. Additionally, PCC was computed between the prediction and the fluorescence density datasets.

#### VSCyto2D

For HEK293T and A549, segmentation of fluorescence density images as well as virtual staining prediction is performed with the ‘nuclei’ (nuclei) and ‘cyto3’ (cells) models in Cellpose. For BJ-5ta, the ‘nuclei’ model in Cellpose is used for nuclei segmentation, and a fine-tuned ‘cyto3’ model (‘CP_20240530_060854‘) was used for cell segmentation. The nuclei segmentation target is corrected by a human annotator. Pearson correlation coefficient (PCC) is computed between the virtual staining prediction and fluorescence density images. Average precision at IoU threshold of 0.5 (AP@0.5) is computed between segmentation masks generated from virtual staining images and segmentation masks generated from fluorescence density images.

For iNeuron, the soma segmentation is performed with the ‘cyto3’ model in Cellpose. The neurites are traced from Calcein fluorescence or virtual staining with scikit-image (54), by multiplying the image with its Meijering-ridge-filtered (55) signal, applying Otsu thresholding (56), removing small objects, and skeletonizing (57). The total neurite length in each FOV is approximated by taking the sum of foreground pixels in the neurite traces. To count the number of neurites connected to each soma, the following steps were taken: 1) the soma foreground mask is first subtracted from the neurite traces, 2) the soma labels are then expanded for 6 pixels (∼2 µm) without overlapping, 3) the number of neurite segments intersecting with these expanded rings that are more than 100 pixels long, are counted as belonging to the respective soma instances.

#### Cellpose Segmentation Parameters

The following parameters are used for segmenting the experimental fluorescence and virtual staining to evaluate the respective models.

##### VSCyto3D

|  |  |  |
| --- | --- | --- |
| **Target** | Nuclei | Cell body |
| **Cellpose model** | 'CP_20220902_NuclFL' | 'cyto3' |
| **Diameter** | 0.0 | 200.0 |
| **Flow Threshold** | 0.4 | 0.4 |
| **Cell probability threshold** | 0.0 | 0.4 |
| **min_size** | 15 | 15 |

##### VSNeuromast

|  |  |  |
| --- | --- | --- |
| **Target** | Nuclei | Cell body |
| **Cellpose model** | 'cellpose_Slices_decon_nuclei_nuclei_v7_2023_06_28_16_54' | 'cellpose_2Chan_scratch_2024_04_30_11_12_00' |
| **Diameter** | 60.0 | 65.0 |
| **Flow Threshold** | 0.0 | 0.4 |
| **Cell probability threshold** | 0.0 | 0.0 |
| **min_size** | 8000 | 8000 |

##### VSCyto2D

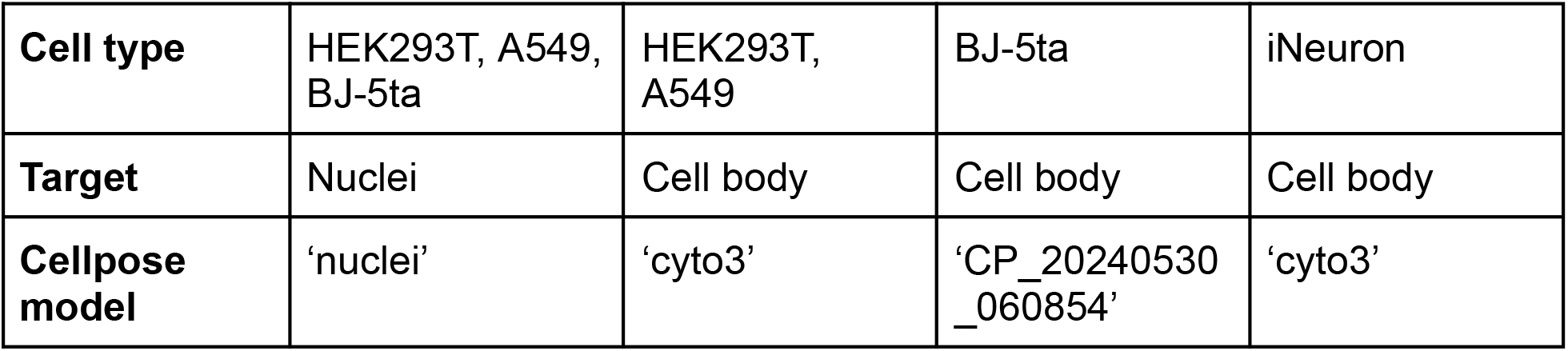
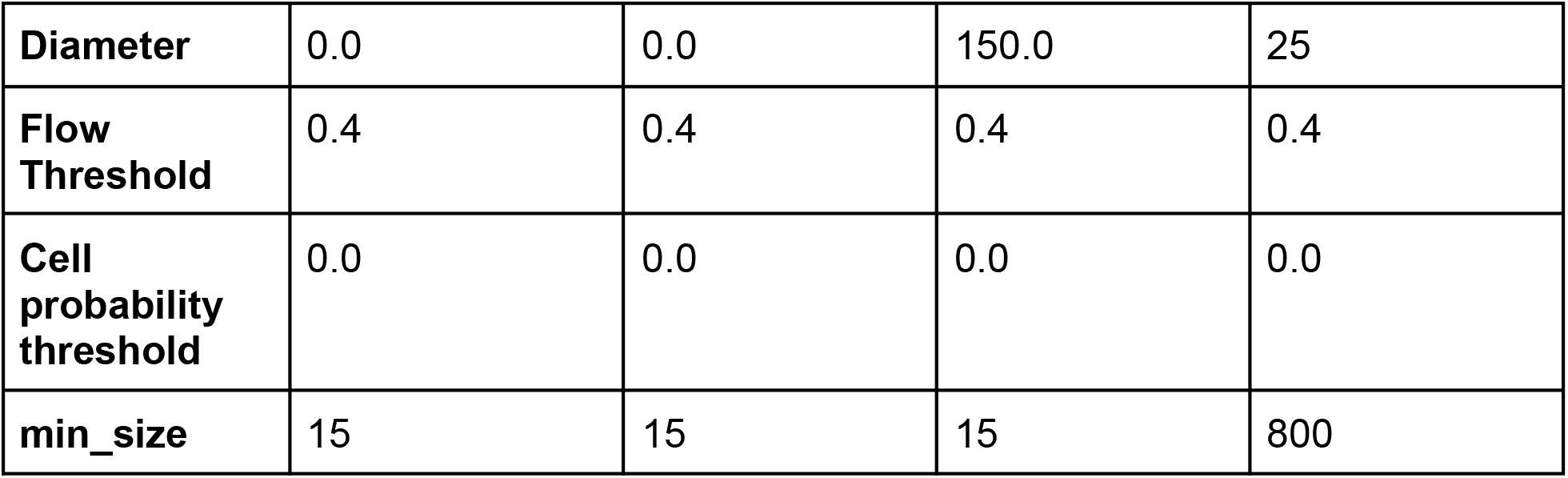

### Model visualization

#### Principal component analysis of learned features

Each XY pixel in the output of a convolutional stage is treated as a sample with channel dimensions and decomposed into 8 principal components. The top-3 principal components are normalized individually and rendered as RGB values for visualization.

## Code and Model Availability

The virtual staining pipeline is implemented as part of an open-source Python package for single-cell phenotyping, named <u>VisCy</u> (a contraction of words ‘vision’ and ‘cell’). We use PyTorch (58) and PyTorch Lightning (59) as the training framework, MONAI (44) for data augmentation, and PyTorch Image Models (timm) (60) for building blocks. Additionally, we implement custom I/O modules for reading and writing OME-Zarr stores during training and inference. The models referenced in the methods above are shared with releases and on the wiki of VisCy via GitHub (34).

## Data Availability

Illustrative test datasets are accessible from scripts in released versions of VisCy. We will share the training datasets via public archive as the review of this work progresses. Some of the datasets are already available from <u>our server</u>.

## Acknowledgments

Some of the computational experiments reported in this paper are informed by the discussions with participants of the advanced research course on deep learning at Marine Biological Lab (DL@MBL), Woods Hole. S. B. M. thanks fellow faculty Anna Kreshuk, EMBL, Heidelberg and Alex Krull, University of Birmingham, for critical discussion of strategies. E. H. thanks Ashesh, Human Technopole, Milan for pair coding image translation models during DL@MBL 2023. We thank Janie Byrum and George Bell, Chan Zuckerberg Biohub for their help with cell culture. We thank CZ Biohub’s

Scientific Computing team for enabling access to the high performance computing cluster. We thank Priscilla Chan and Mark Zuckerberg for supporting the CZ Biohub.

## Author contributions

Z. L., E. H-M., S. P., J. R., I. I., H. W., T. L., A. B., R.M., C. L., A. J., and S. B. M. contributed imaging data for training and testing the models. Z. L., E. H-M., S. P., C. F., and S. B. M. contributed to the development of the code for training models. Z. L. and E. H-M trained and evaluated models reported in this paper with guidance from S. B. M. R. A., M. L., C. A., A. J., and S.B.M. supervised the research and informed the evaluation of the models. Z. L., E. H-M, and S. B. M. wrote the paper with critical contributions from all co-authors.

## Supplementary Materials

**Figure S1:**
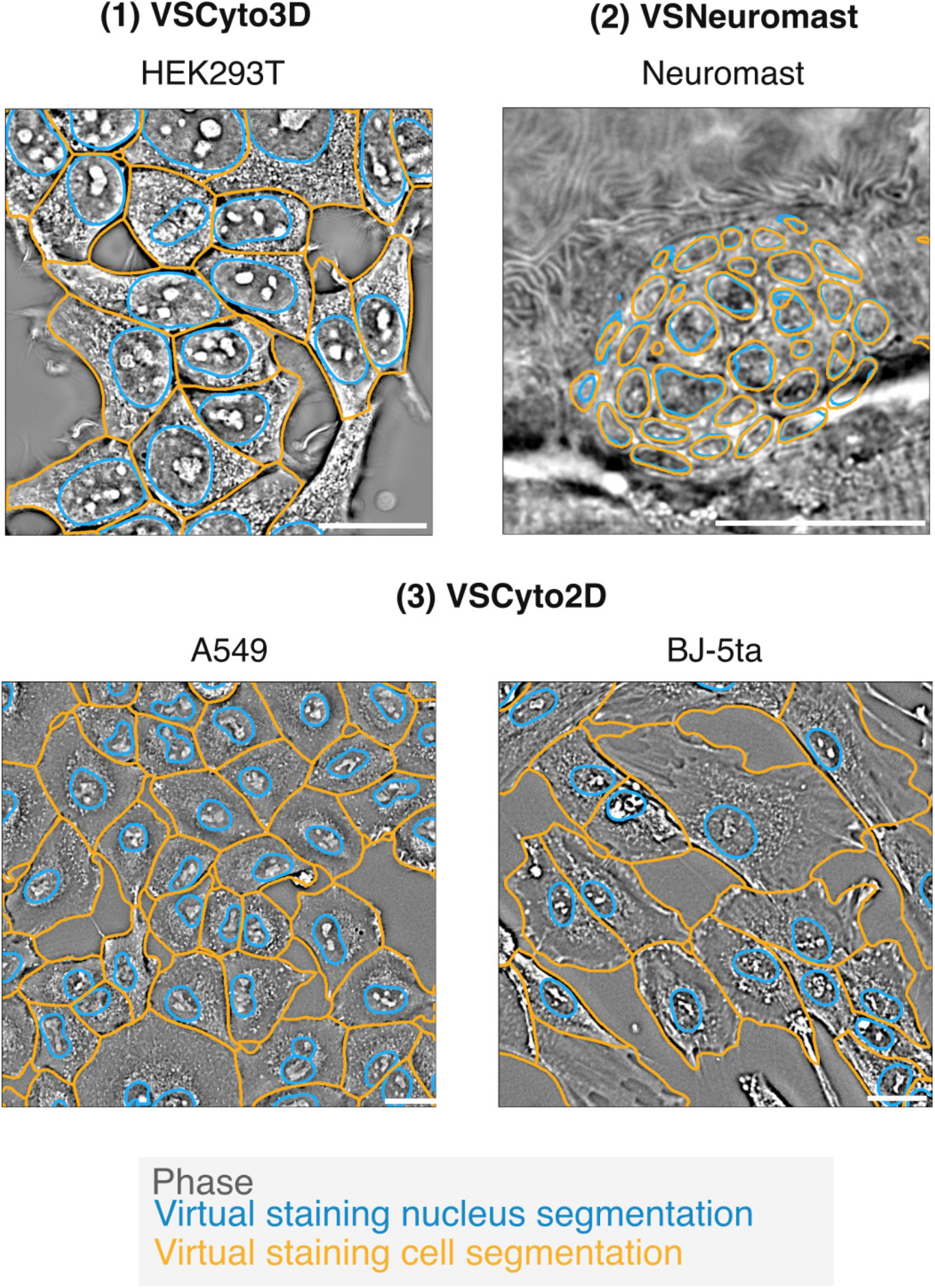
Examples of nuclei and cell segmentation from virtual stained images overlaid on the corresponding phase images.

**Figure S2:**
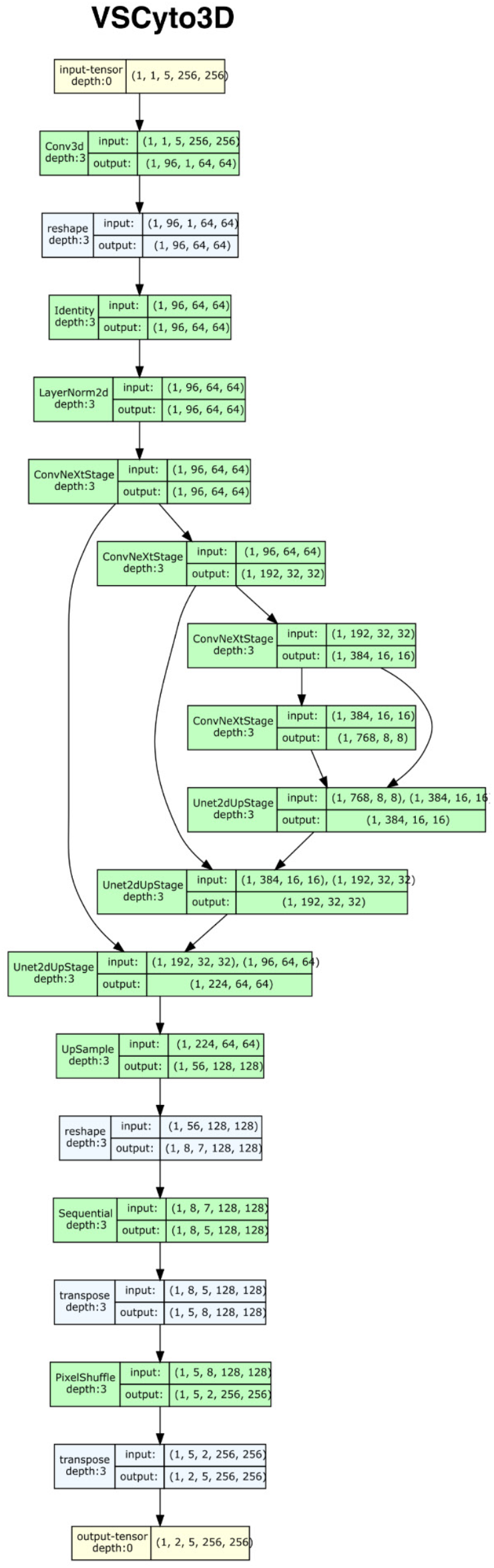
The model architecture used to train VSCyto3D

**Figure S3:**
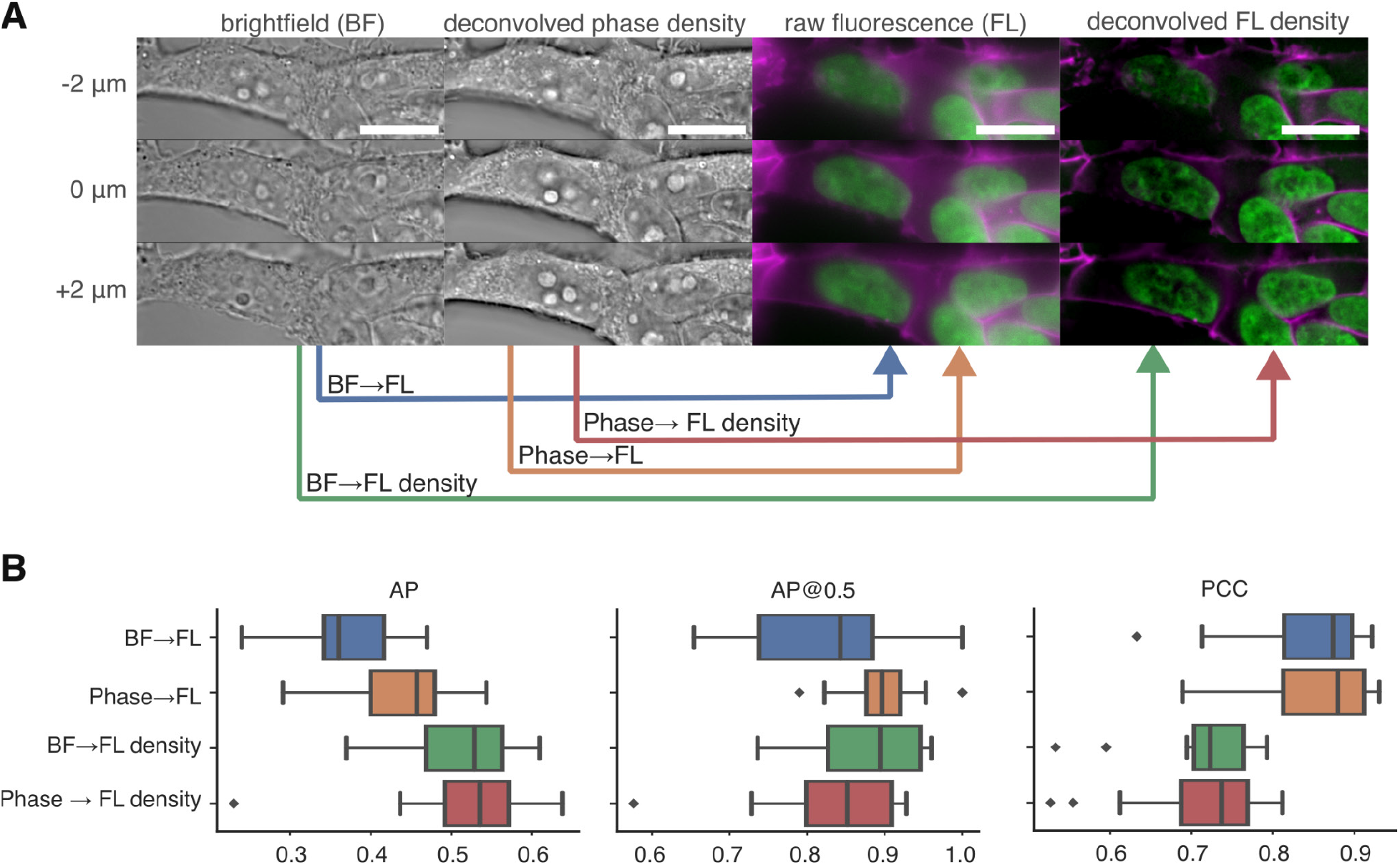
Deconvolution improves contrast of fine features for virtual staining. **(A)** Comparison of contrast in brightfield (BF) and fluorescence (FL) images with the corresponding deconvolution of the phase density and fluorescence density. We trained four virtual staining models that translate between raw and deconvolved versions of label-free and fluorescence contrasts (indicated by arrows). Scale bars: 10 µm. **(B)** The average precision (AP) and average precision at IoU =0.5 (AP@0.5) for nuclei segmented from experimentally and virtual stained images are shown. Virtually stained images were predicted with four models indicated on the y-axis. Instance segmentations of experimentally stained nuclei were proofread manually. Deconvolution of BF and FL volumes leads to more accurate segmentation of nuclei. We also assess how the phase and fluorescence density, deconvolved from brightfield (BF) and fluorescence (FL) volumes, respectively, affect the Pearson cross-correlation (PCC).

**Figure S4:**
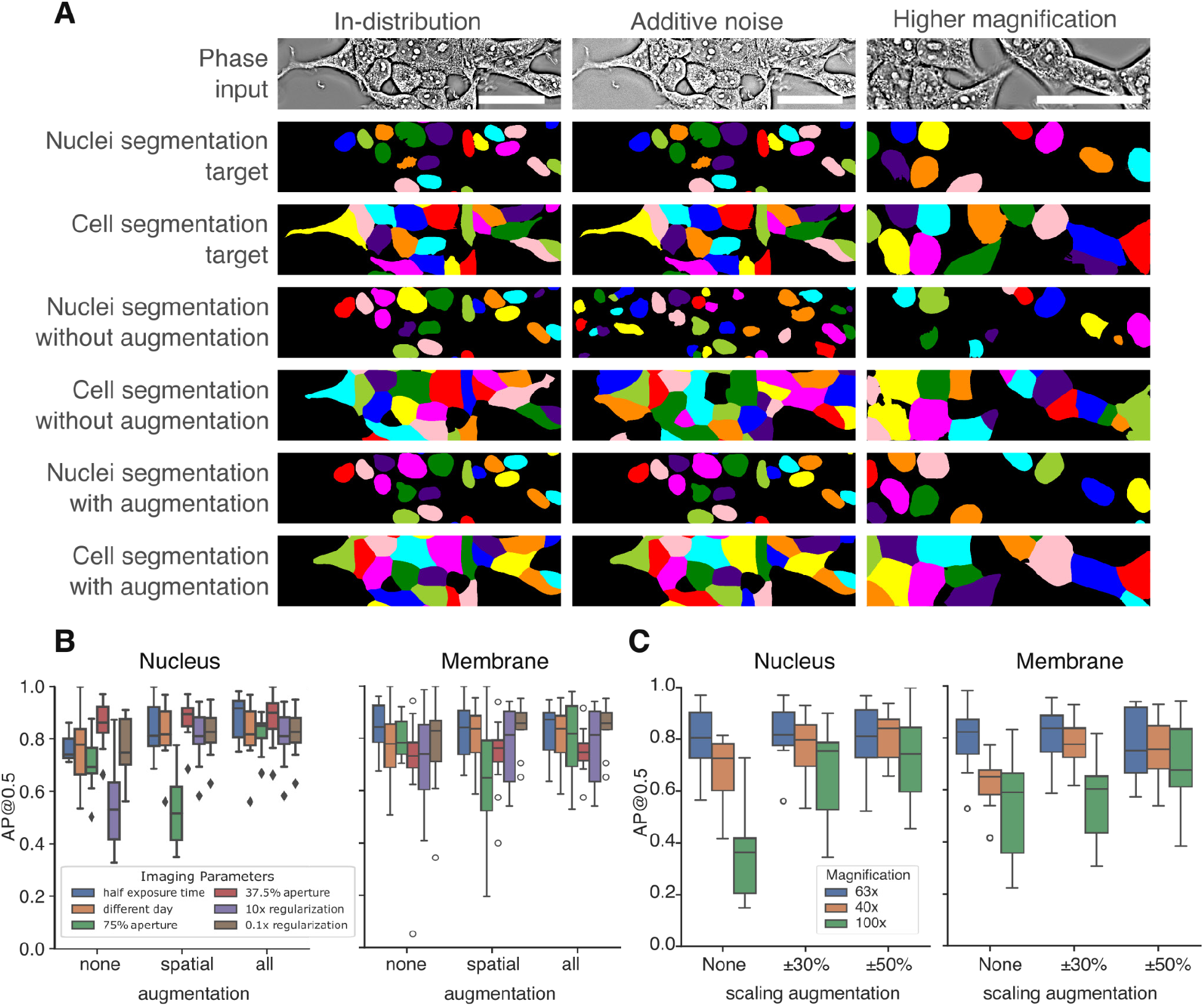
Augmentation-induced robustness in virtual staining improves segmentation of nuclei and cells **(A)** Segmentations from virtual staining predictions of VSCyto3D models trained without augmentation are inaccurate when noise is added to the phase image or when phase image at a different magnification is used for inference. The segmentations are more reliable when using augmentations as described in <u>Methods</u>. Scale bars: 50 µm. **(B)** Test FOVs were acquired with diverse imaging parameters (see the legend). Virtual staining models trained without augmentation (group: none), with spatial augmentation (group: spatial), and with spatial and intensity augmentations (group: all) were used to virtually stain nuclei and membranes in test FOVs. We compare the AP@0.5 between instance segmentations from virtually stained nuclei and proofread instance segmentations from experimentally acquired nuclei images. The solid lines in the box plot indicate the median and interquartile ranges of AP@0.5 across all FOVs in the test set. The predictions of models become invariant to changes in the imaging parameters as more data augmentations are used during training. AP@0.5 for membrane segmentation also indicates that the augmentations make the membrane predictions invariant to imaging parameters. **(C)** AP@0.5 for test FOVs acquired at the same (63x), higher (100x), and lower (40x) magnifications relative to the training dataset, when no scaling augmentation (group: none), and increasing scaling augmentation (groups: ±30%, ±50%) is used when training models for joint virtual staining of nuclei and membrane. The metrics for both nuclei and membranes indicate that the scaling augmentations make the predictions equivariant to magnification.

**Figure S5:**
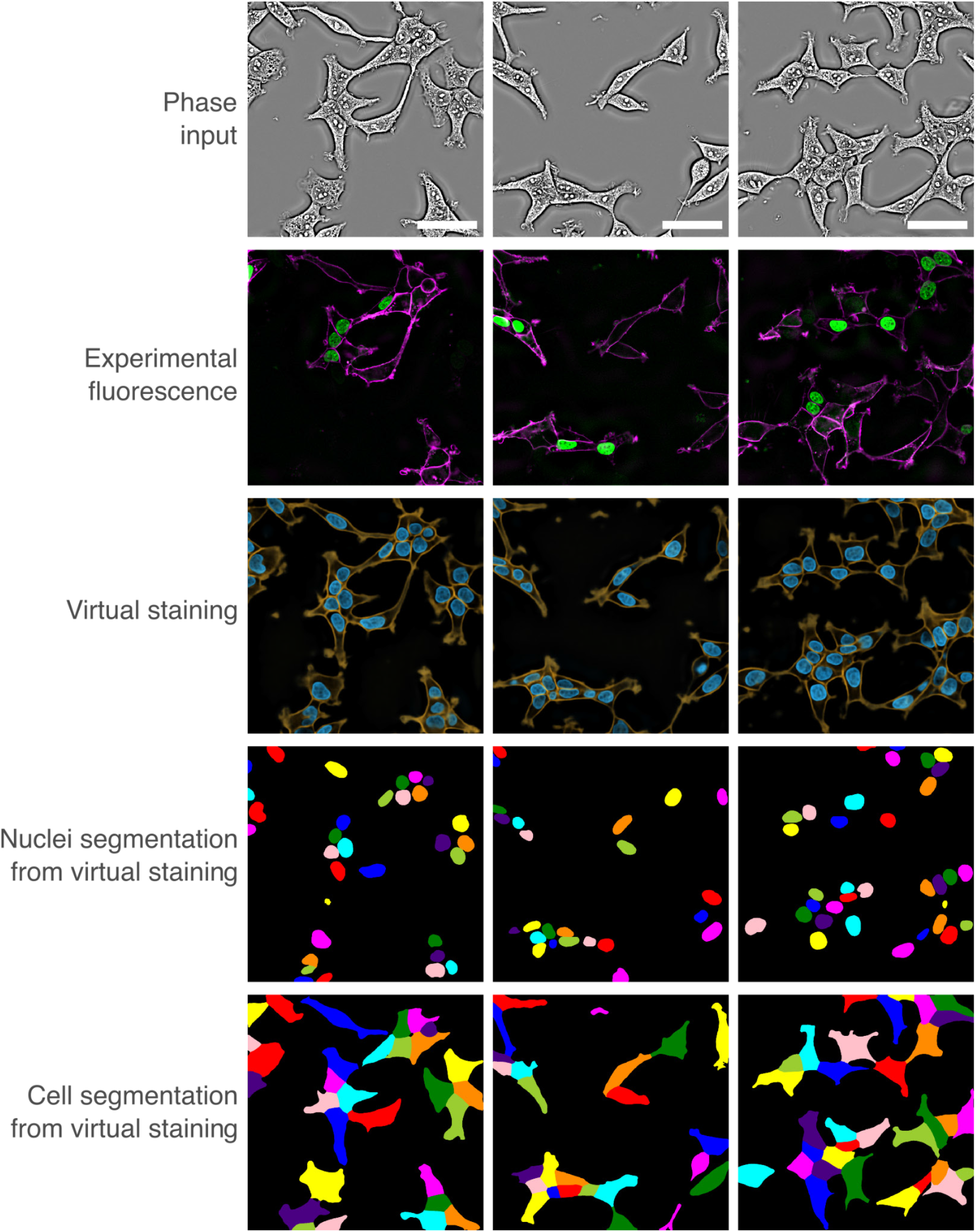
Virtual staining from phase rescues missing fluorescence labels: Experimentally and virtually stained in HEK293T cells nuclei and membrane for 75% aperture and corresponding segmentations. Scale bars: 50 µm.

**Figure S6:**
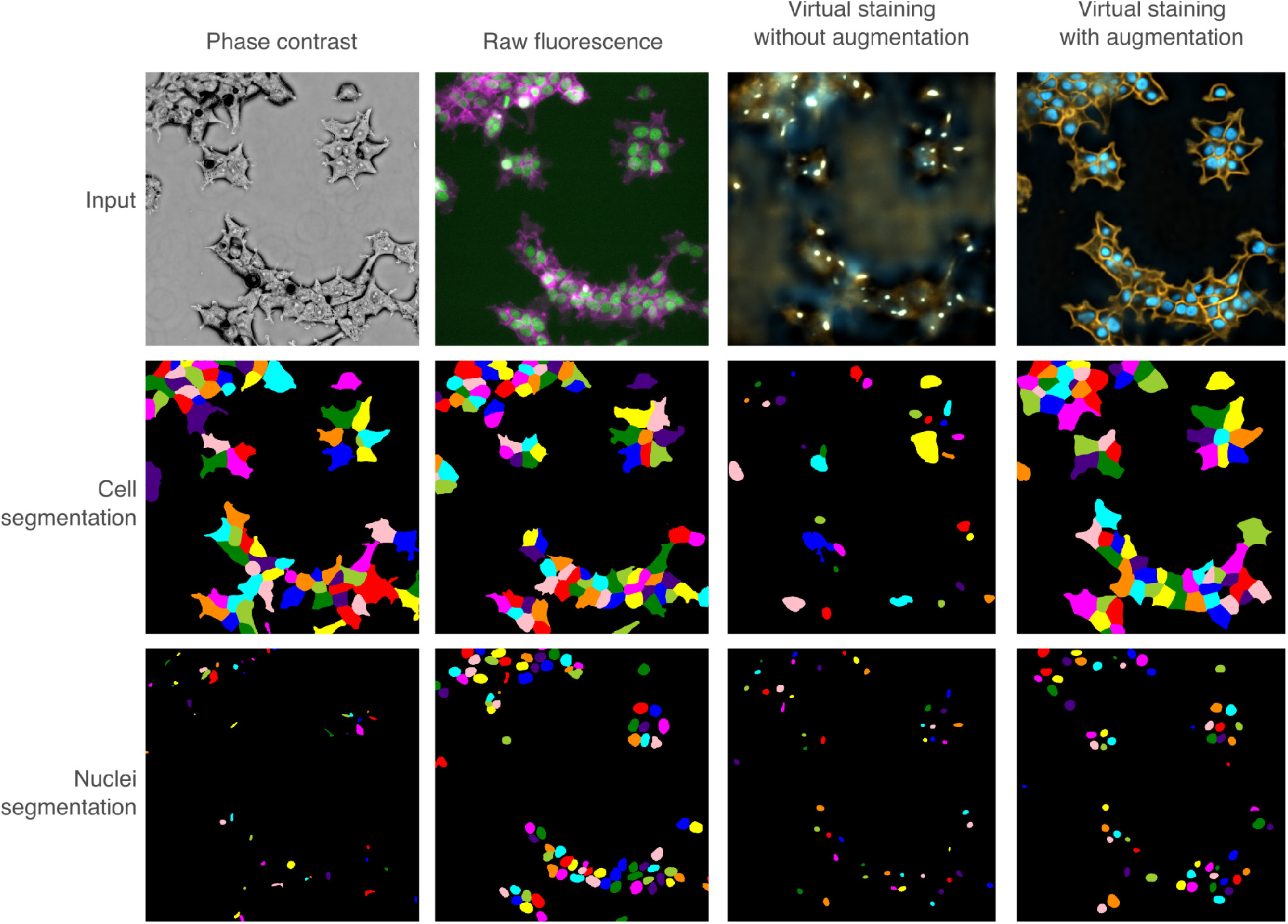
Cell and nuclei segmentation from Zernike phase contrast, raw fluorescence, and virtual staining images. Augmentation improves the virtual staining prediction for segmentation.

**Figure S7:**
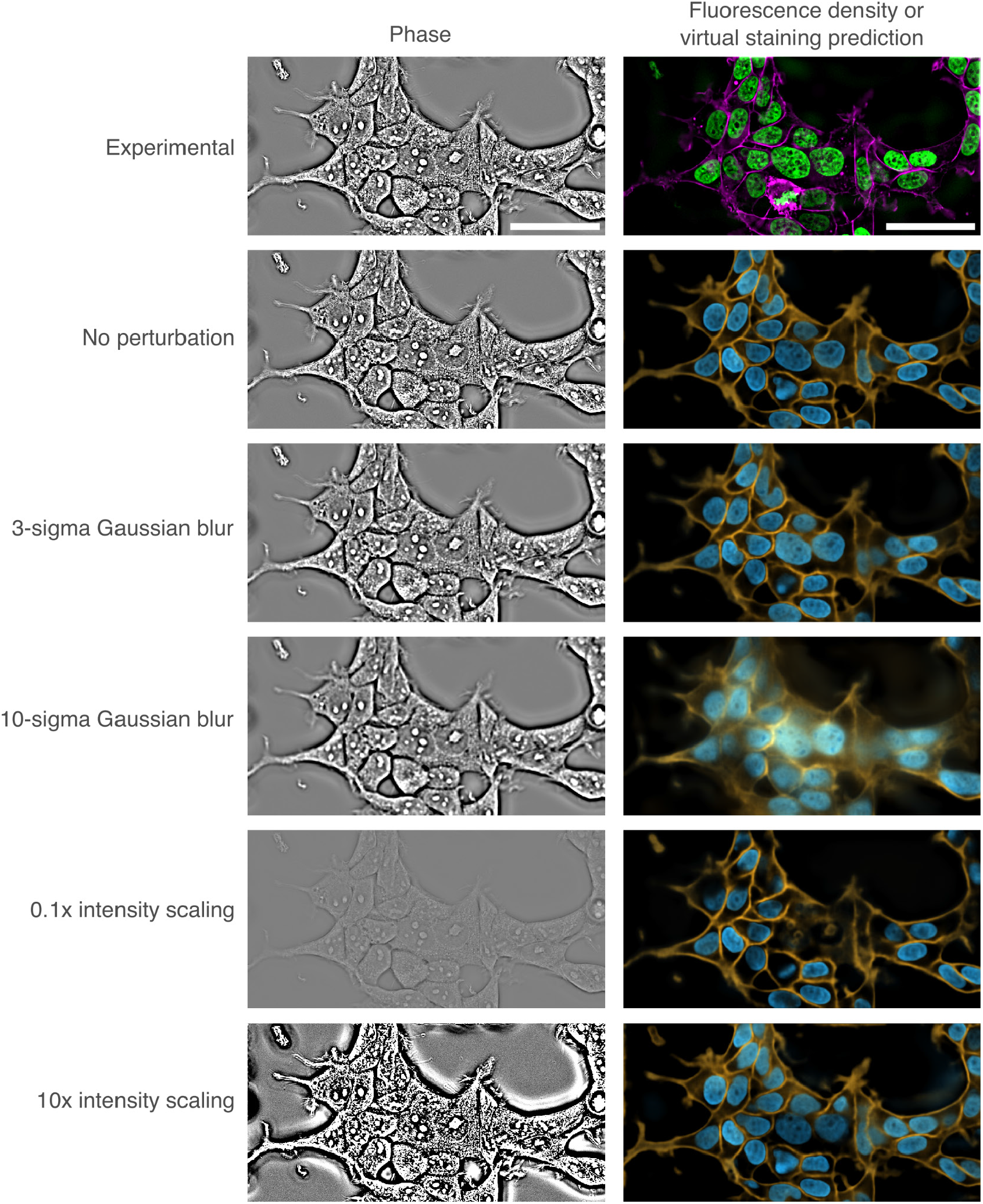
VSCyto3D model’s robustness to perturbations in input: Simulated image perturbations are applied to the input phase image. The virtual staining prediction is robust to a certain range of input perturbation and fails when the deviation is too large (σ=10 pixels). For example, when a very strong Gaussian blur is applied to the input, the model cannot differentiate the in-focus slice from a defocus slice of the imaging volume, and predicts a blurry image. Scale bars: 50 µm.

**Figure S8:**
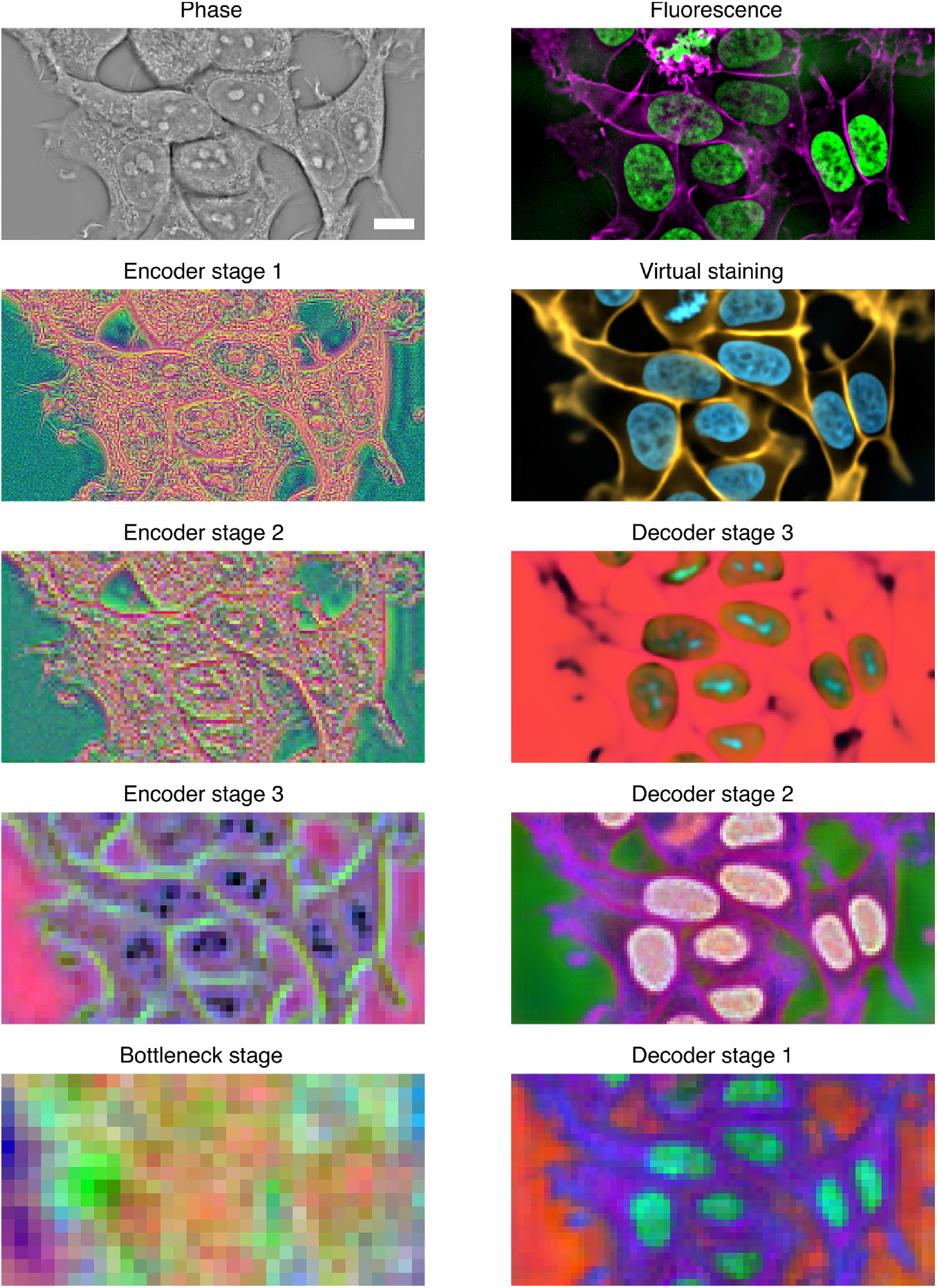
Visualization of features learned by VSCyto3D: Input, prediction, and intermediate feature maps of the 3DVSCyto (Figure 2B ‘deconvolved -> deconvolved’) model trained on HEK293T cells. The first 3 principal components of the feature map from each ConvNext stage are rendered as RGB values for an illustrative input image patch. Scale bar: 10 µm.

**Figure S9:**
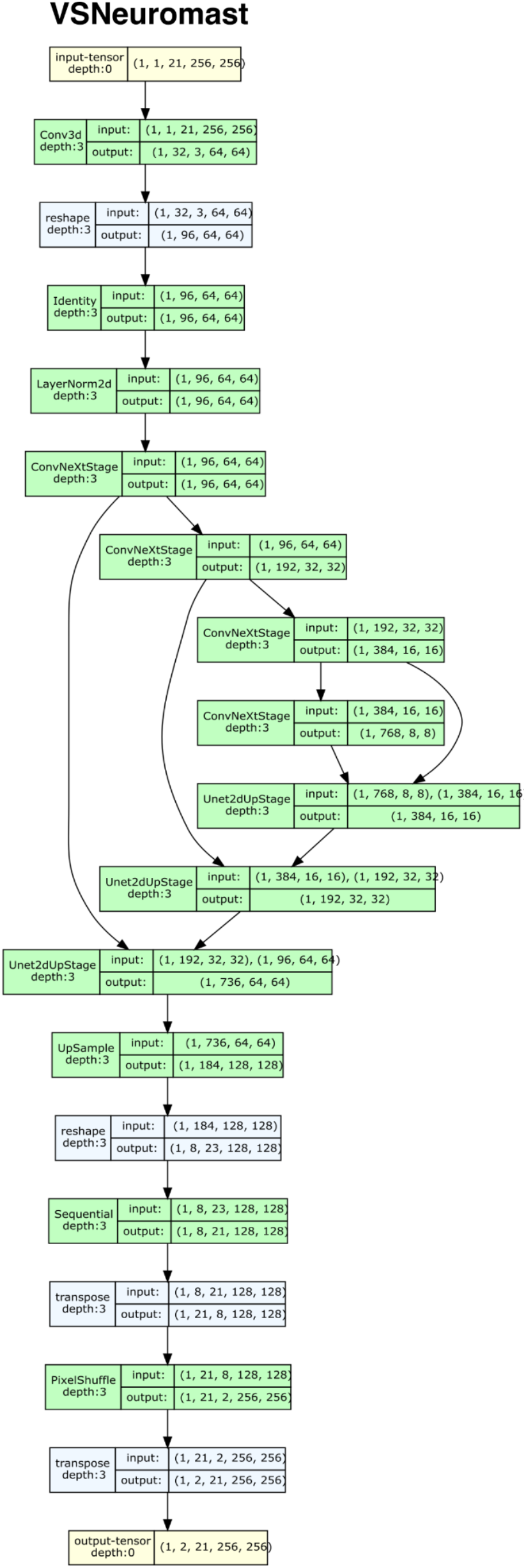
The model architecture used to train VSNeuromast

**Figure S10:**
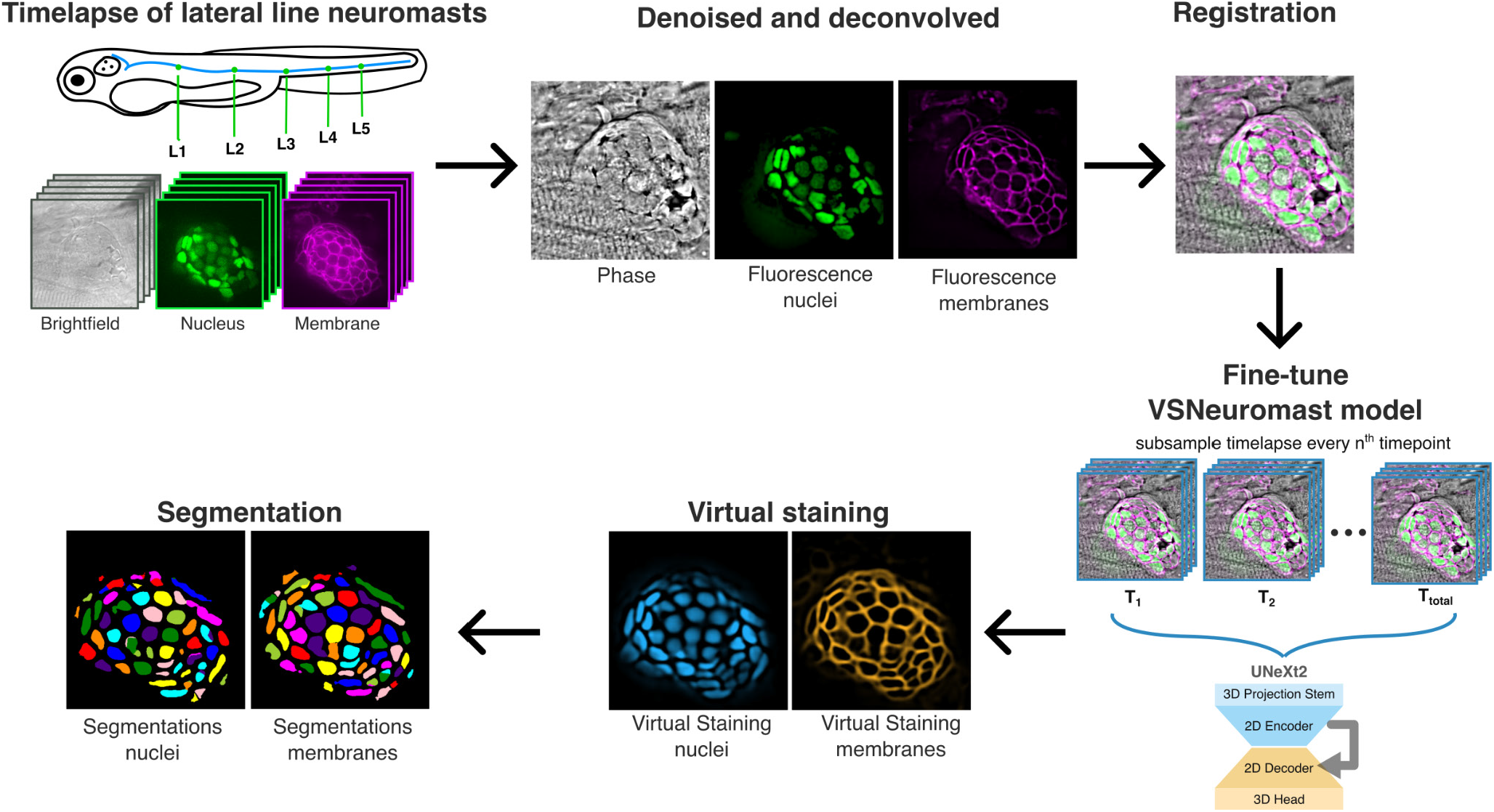
Illustration of the workflow for fine-tuning the VSNeuromast model: The model is fine-tuned to generate accurate predictions and segmentations by subsampling the timelapse and using these time points as training data. Images of neuromasts during preprocessing (denoising, deconvolution, registration) and postprocessing (segmentation) steps are also shown.

**Figure S11:**
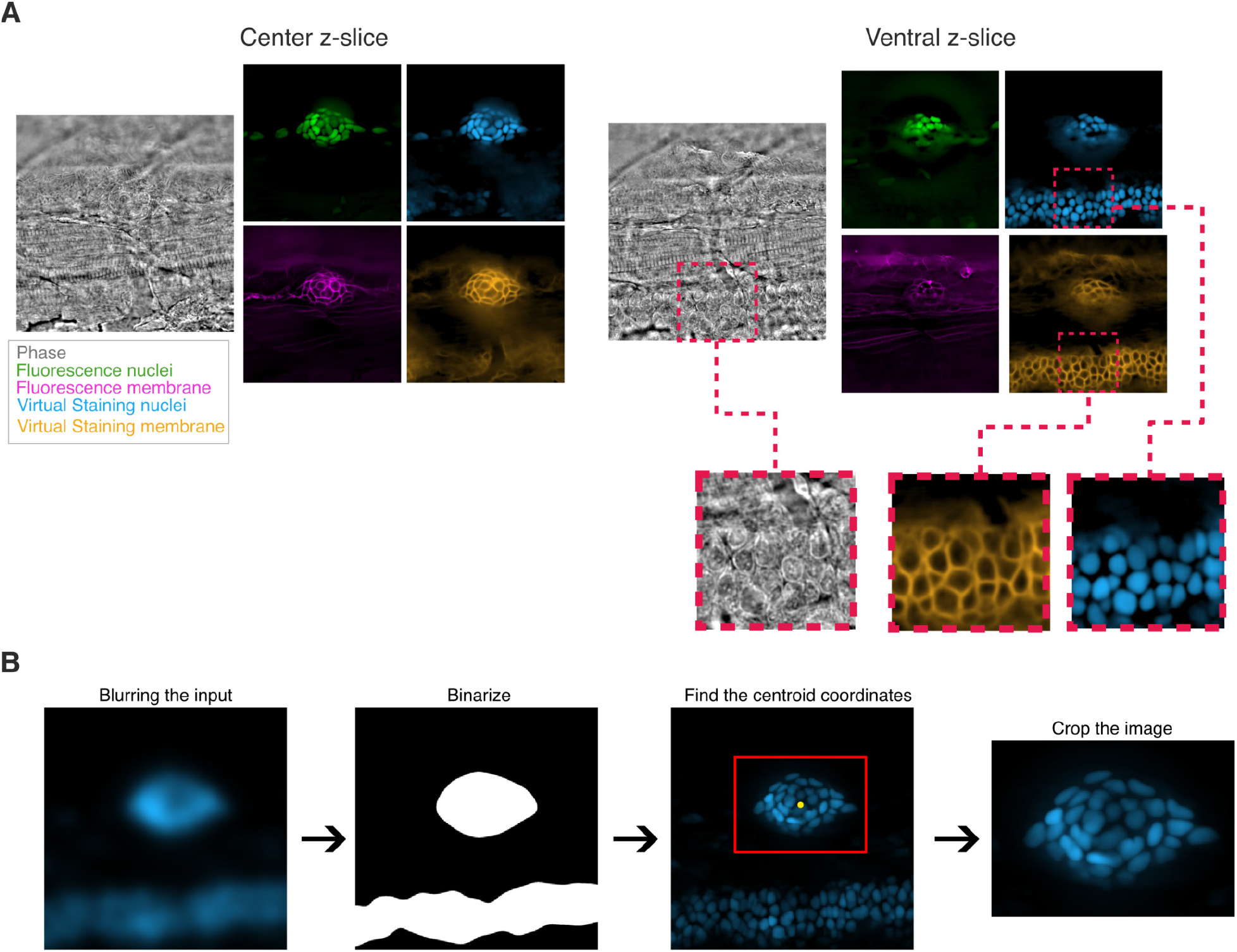
Post-processing is needed to distinguish virtually stained neuromast cells from non-neuromast cells A. Phase, fluorescence and virtual staining pairs of the central and ventral slices depicting how the model generalizes to other cell types with similar morphology. B. Processing pipeline to isolate the neuromasts from the whole FOV. The pipeline is used for generating the instance segmentations and performance metrics.

**Figure S12:**
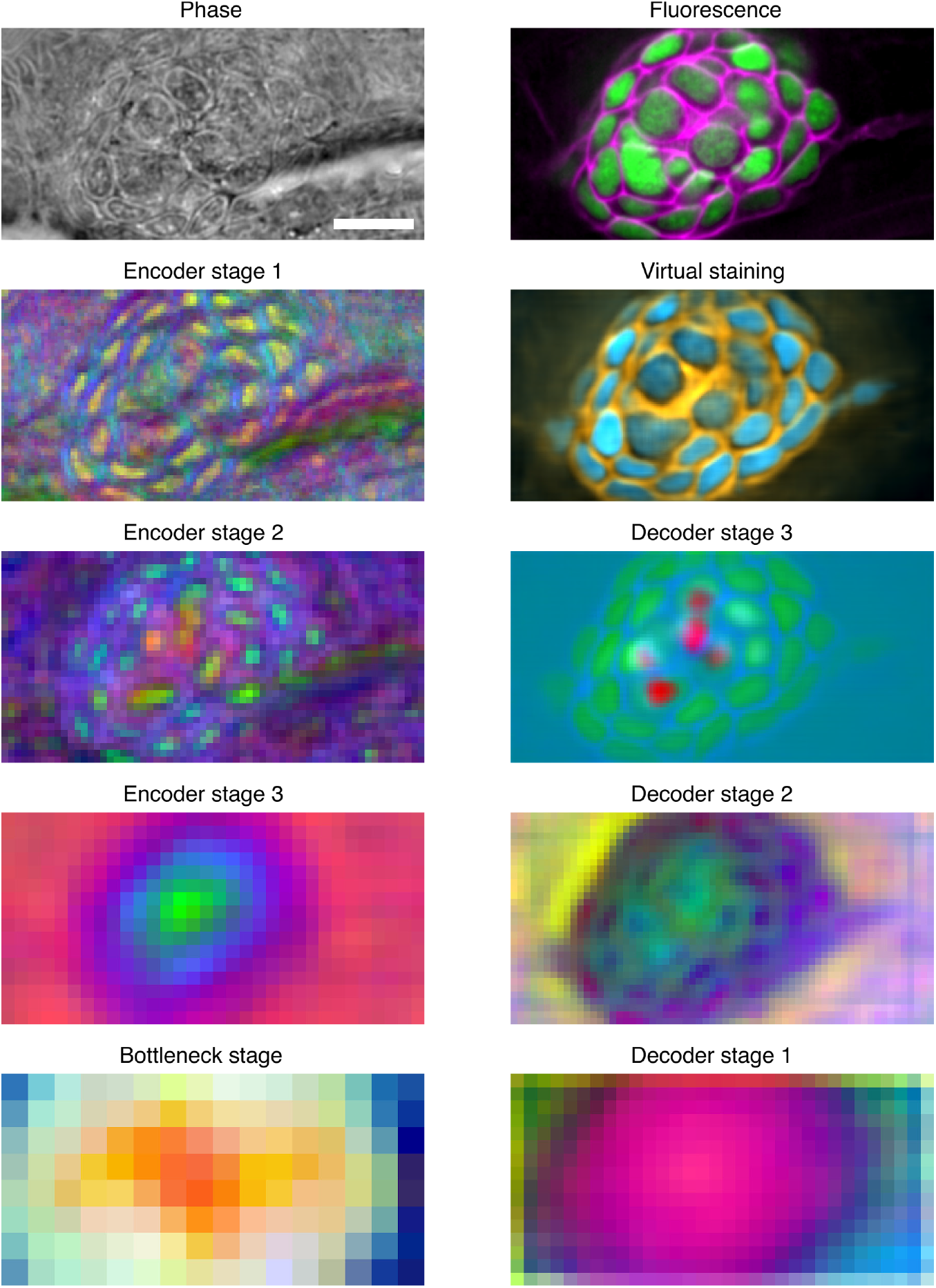
Visualization of **features learned by VSNeuromast**: Input, prediction, and intermediate feature maps of the 3DVSNeuromast model trained on zebrafish neuromasts. The first 3 principal components of the feature map from each ConvNext stage are rendered as RGB values for an illustrative input image patch. Scale bar: 10 µm.

**Figure S13:**
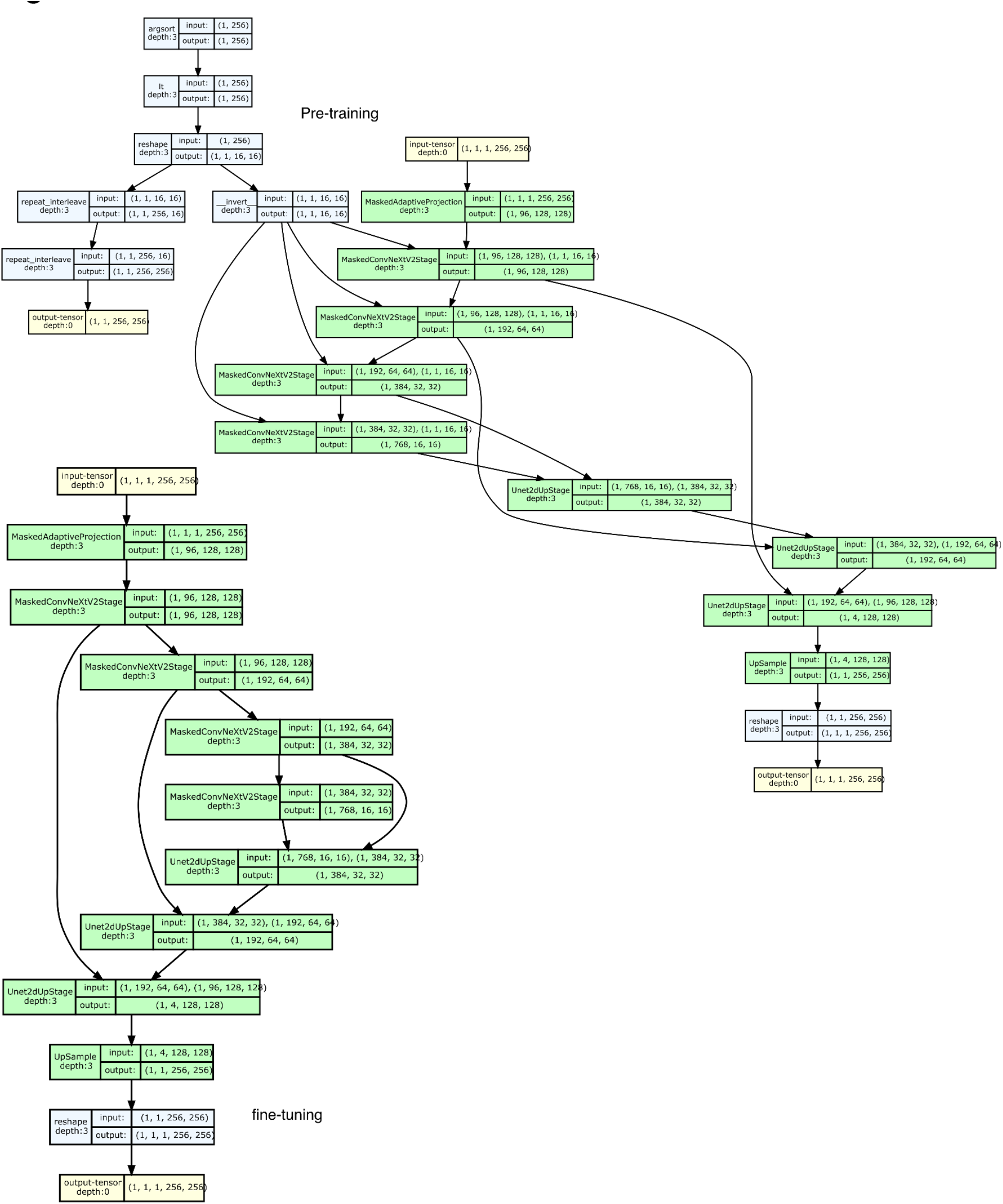
Model architectures used for training VSCyto2D

**Figure S14:**
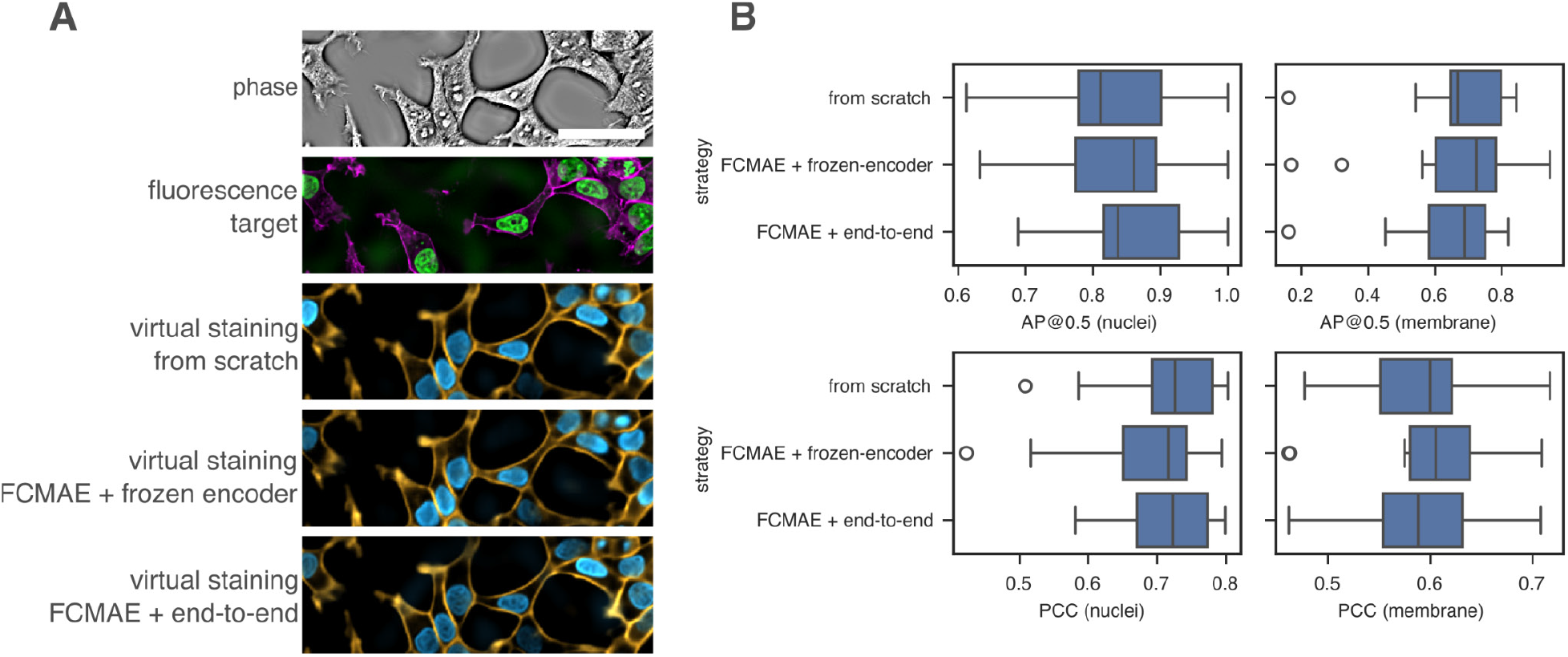
Performance of **virtual staining models of HEK293T cells trained from scratch or with pre-training** **(A)** Virtual staining of nuclei and membrane in HEK293T using models trained from scratch or pre-trained (FCMAE) and then fine-tuned on HEK293T data. pre-training improves the high-frequency features in predictions. Scale bar: 50 µm. **(B)** The pre-training protocol has similar segmentation and regression performance for a single cell type. PCC: Pearson correlation coefficient. AP@0.5: average precision at IoU =0.5.

**Figure S15:**
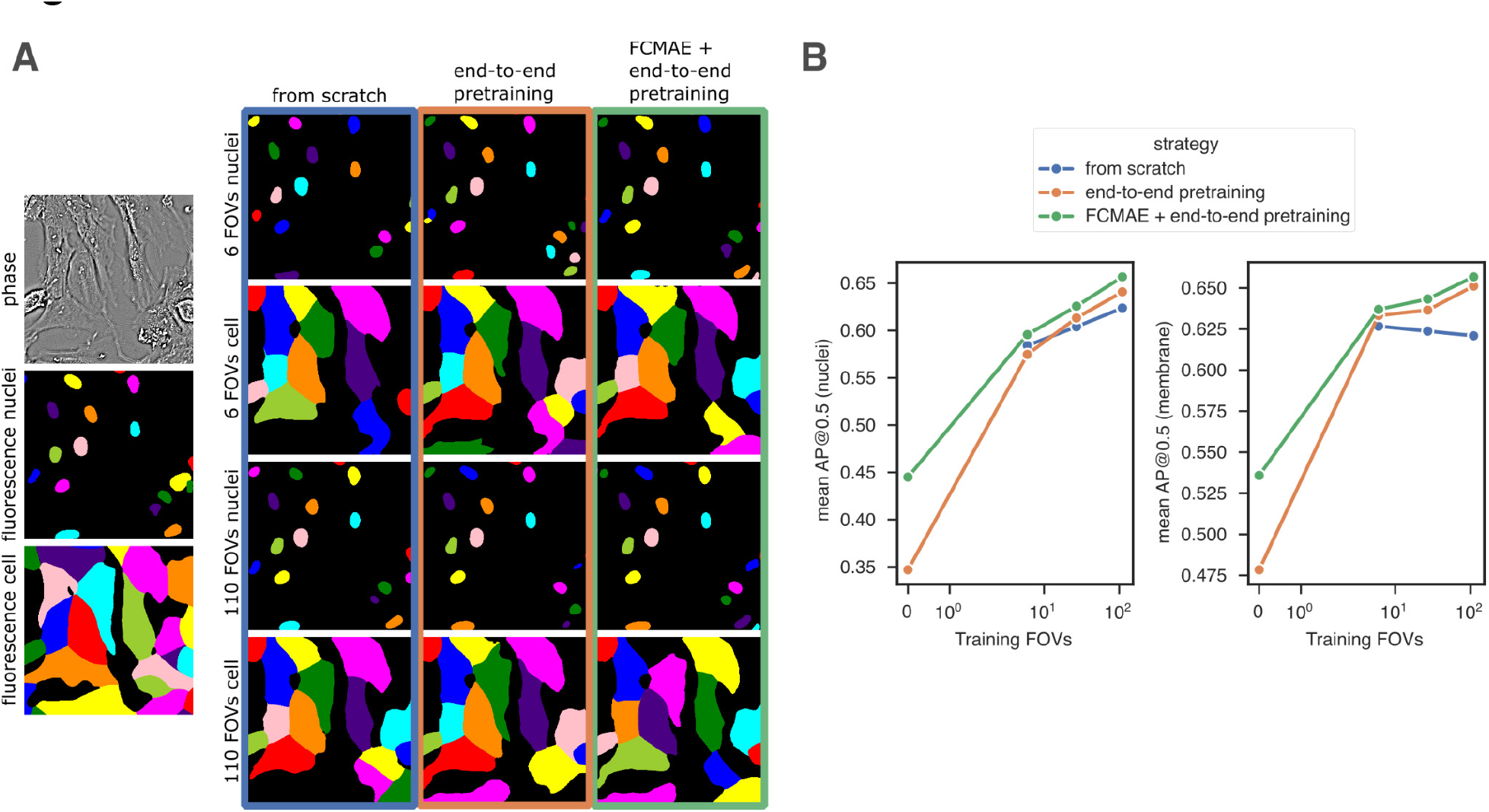
Segmentation performance of virtual staining models of BJ5-ta cells trained from scratch and with pre-training. **(A)** Nuclei and cell segmentation of BJ5-ta cells are shown for the 6-FOVs and 110-FOVs models according to the training strategy described in Figure 4A. **(B)** Segmentation performance from virtual staining in BJ-5ta cells. Pre-training with FCMAE and virtual staining tasks improves segmentation performance. AP@0.5 = average precision at IoU=0.5.

**Figure S16:**
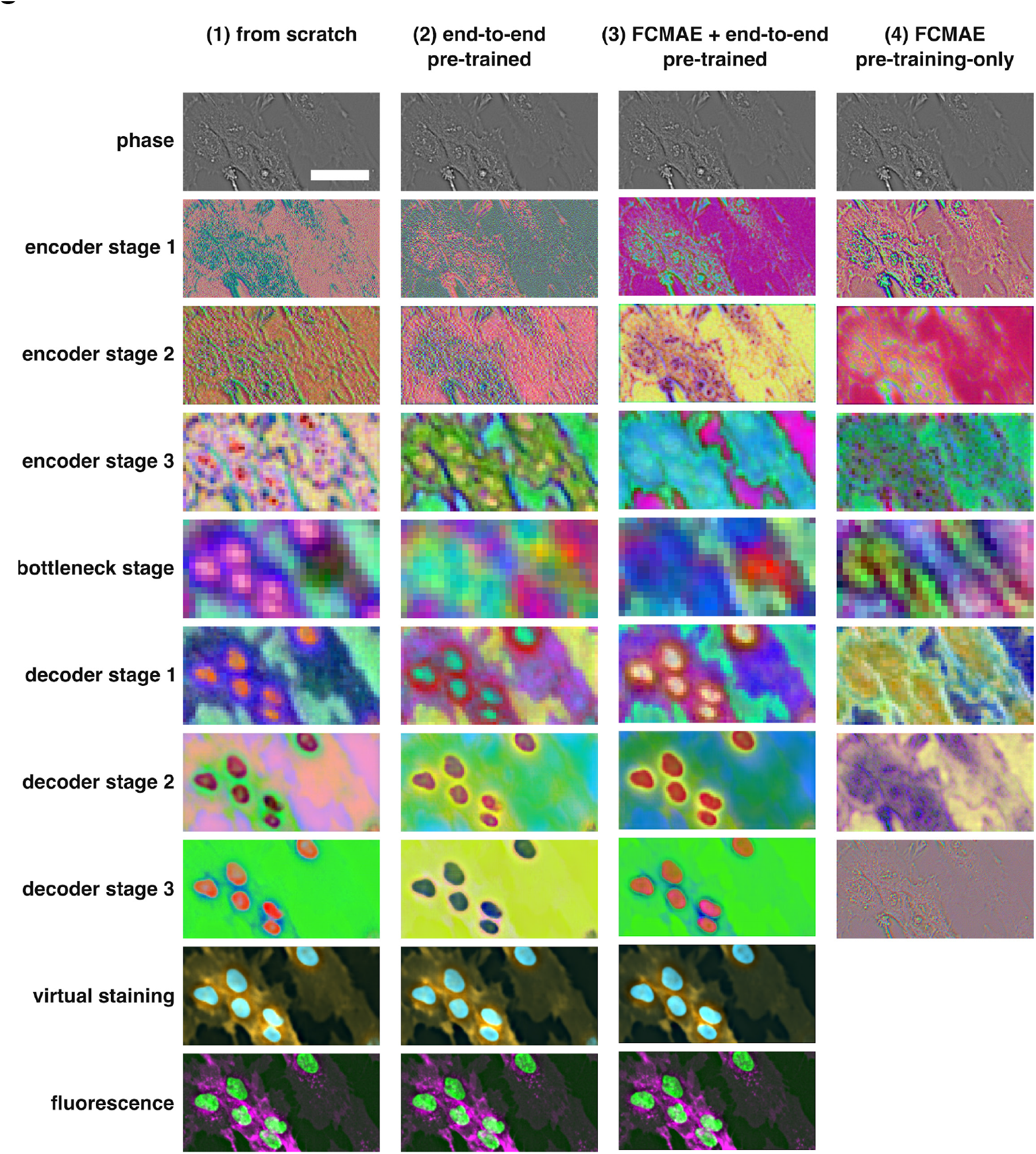
Visualization of features learned by VSCyto2D: Input, prediction, and intermediate feature maps of the 2DVSCyto and FCMAE models. The first 3 principal components of the feature map from each ConvNext stage are rendered as RGB values for an illustrative input image patch. (1) model trained from scratch on BJ-5ta; (2) model pre-trained on virtual staining of HEK-293T and A549, and then fine-tuned on BJ-5ta; (3) model pre-trained with FCMAE and virtual staining of HEK-293T and A549, and then fine-tuned on BJ-5ta; (4) FCMAE model of HEK-293T and A549, not trained for virtual staining. Scale bar: 50 µm.

**Video 1:**
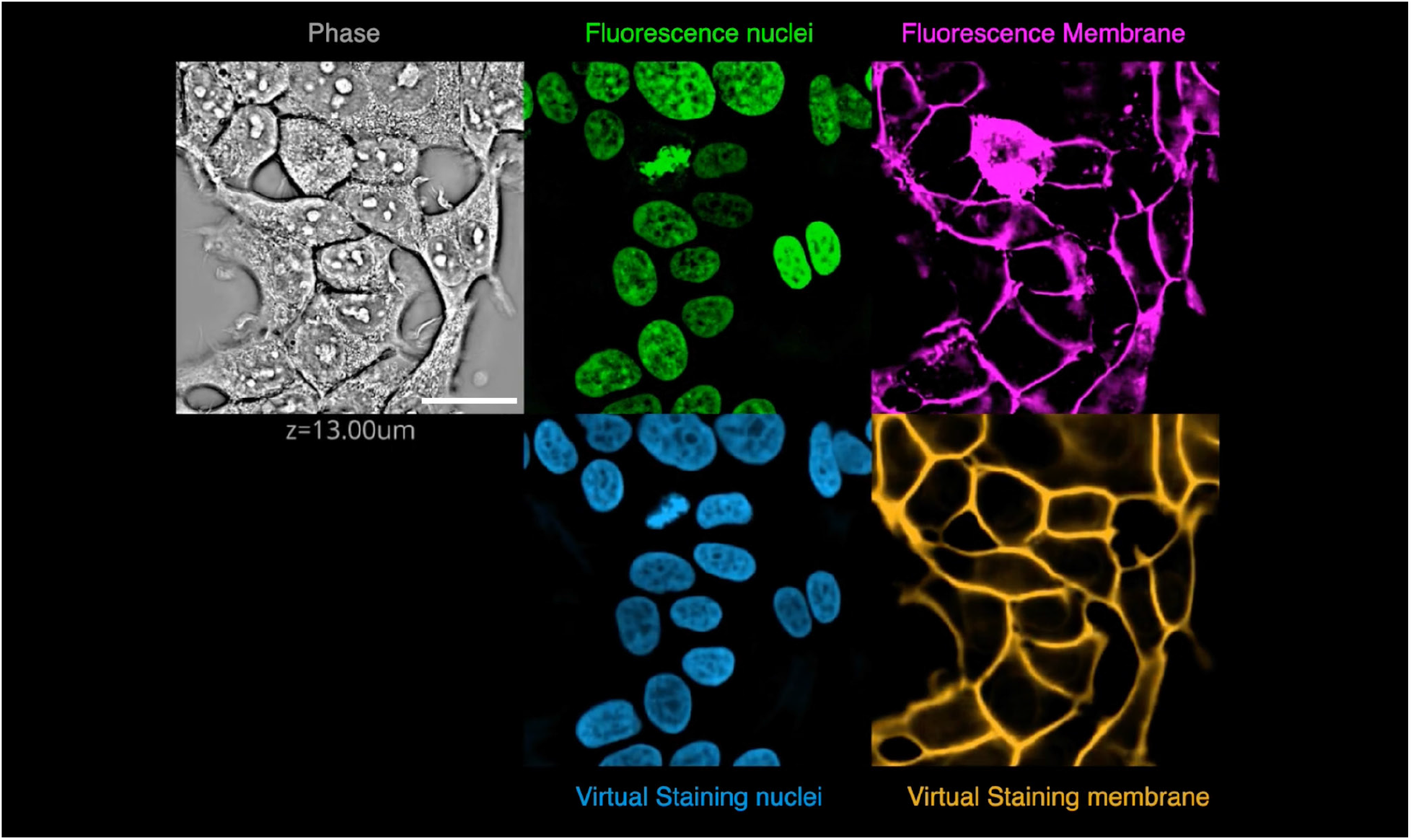
Through-focus movie of HEK293T cells: phase, experimentally stained nuclei (green) and membrane (magenta), virtually stained nuclei and membrane with VSCyto3D. (scale bar 25µm)

**Video 2:**
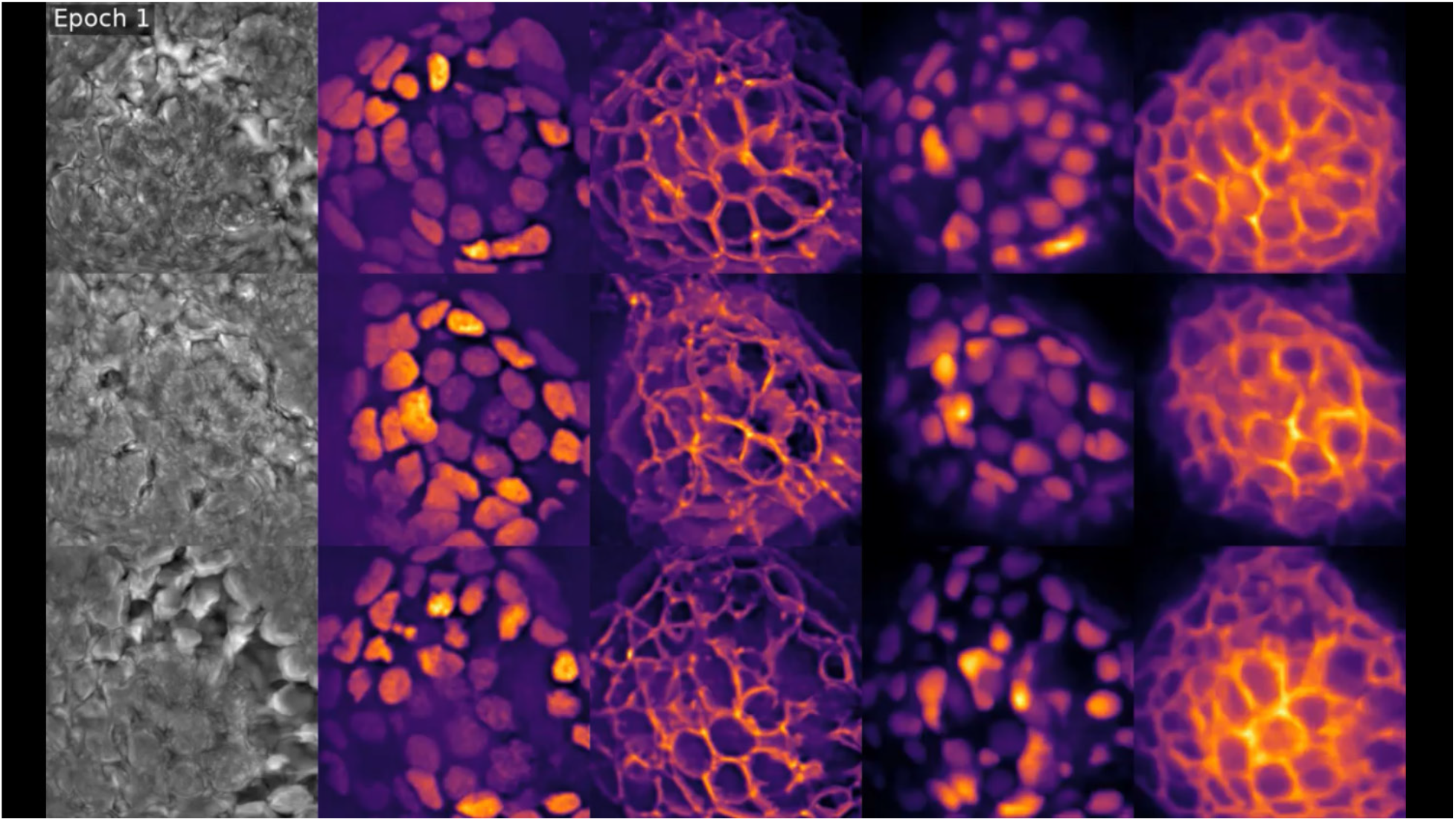
Evolution of neuromast predictions during fine-tuning. The first three columns depict the 2D source (phase) and target (nuclei and membrane) pairs of three different fields of view (FOVs) from the validation dataset. The last two columns feature the virtual staining predictions of nuclei and membrane respectively.

**Video 3:**
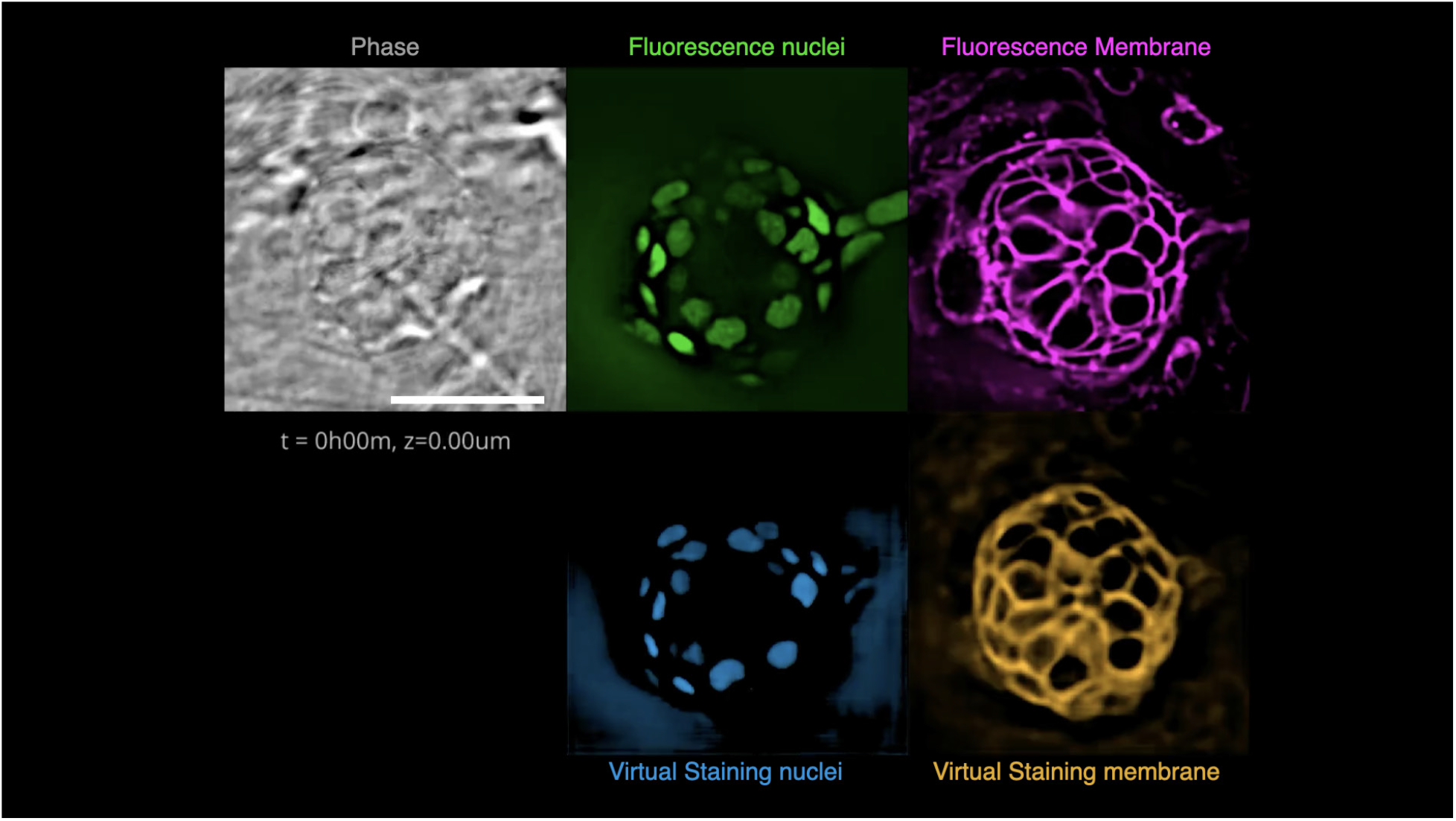
Axial and temporal fly-through of fluorescence and virtually stained landmarks. Displayed are the phase image in gray, fluorescence nuclei in green, and membrane in magenta, along with virtually stained nuclei in blue and membrane in orange, predicted using the fine-tuned VSNeuromast. (Scale bar: 25µm)

**Video 4:**
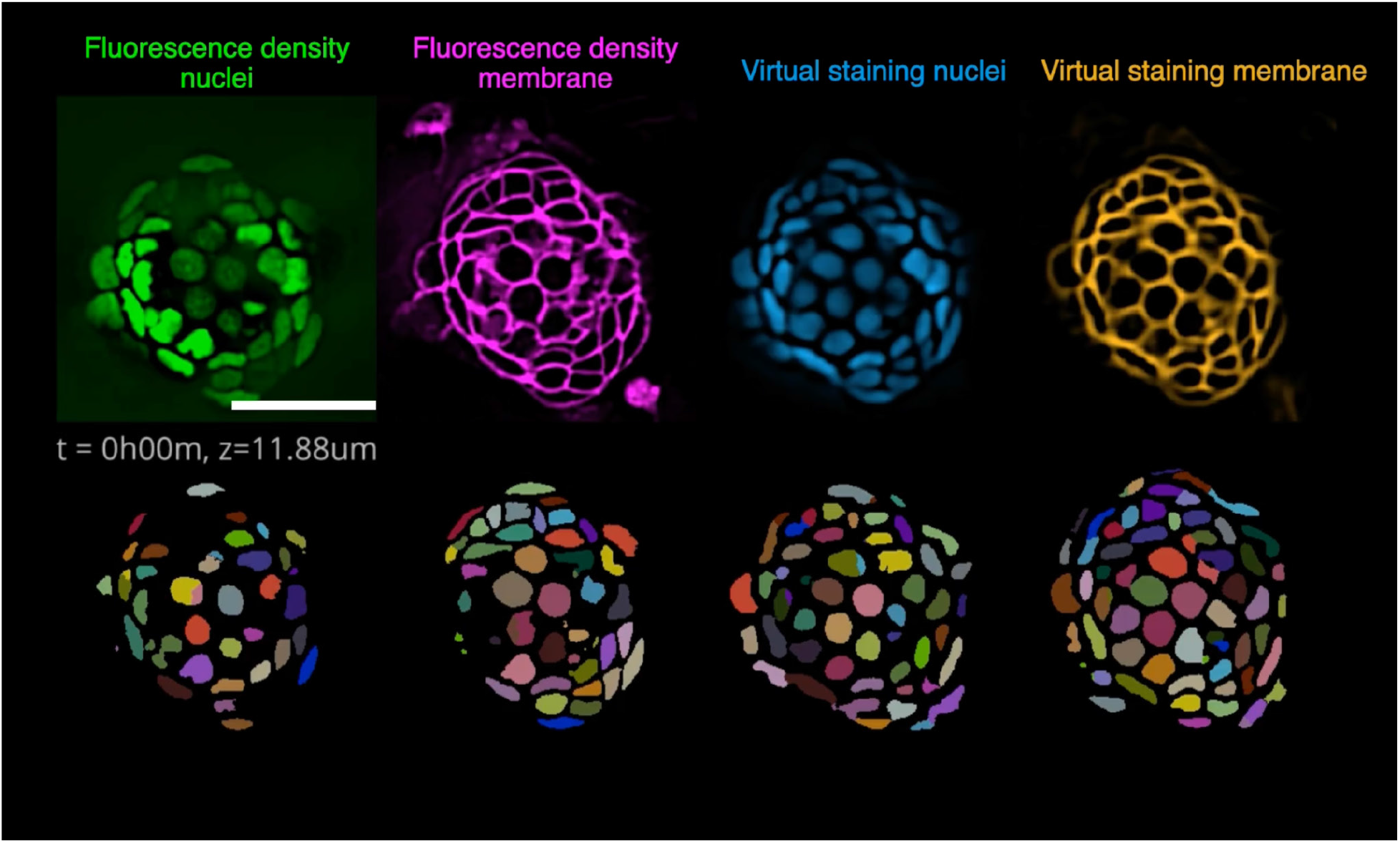
Comparison of neuromast nuclei and membrane fluorescence density and virtual staining 3D segmentations over time. The video shows the 3D instance segmentations at t=0 using the fine-tuned Cellpose model for nuclei and membrane respectively applied to both the fluorescence density and virtual stained and plays over time at the middle z-plane of the neuromast. Virtual staining rescues the uneven expression of nuclei and segments allowing for better segmentation (Scale bar= 25µm)

**Video 5:**
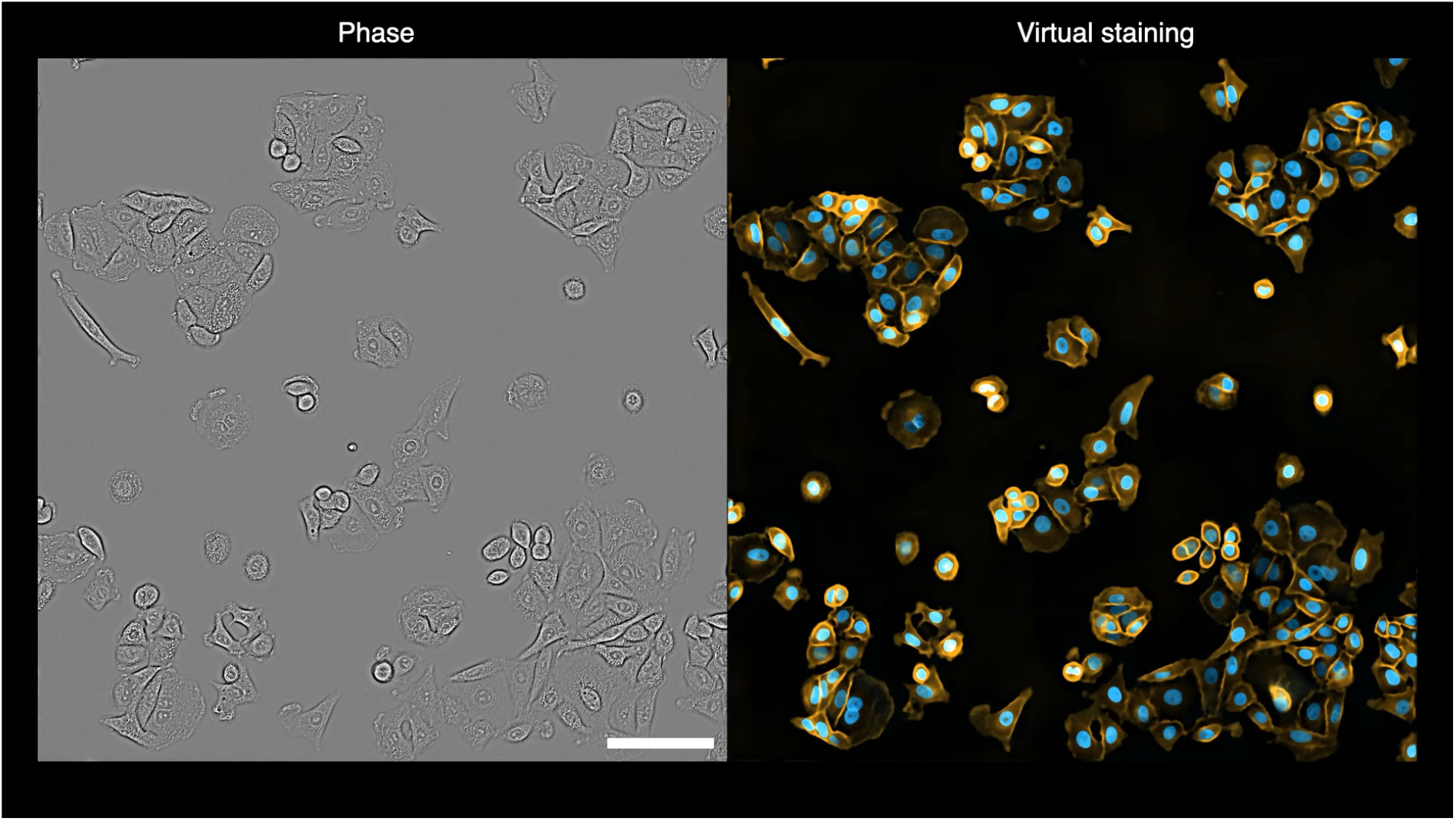
Phase time-lapse images virtually stained with VSCyto2D. Blue: nuclei; Orange: membrane. Every 30 minutes for 24 hours. Scale bar: 100 µm.

**Video 6:**
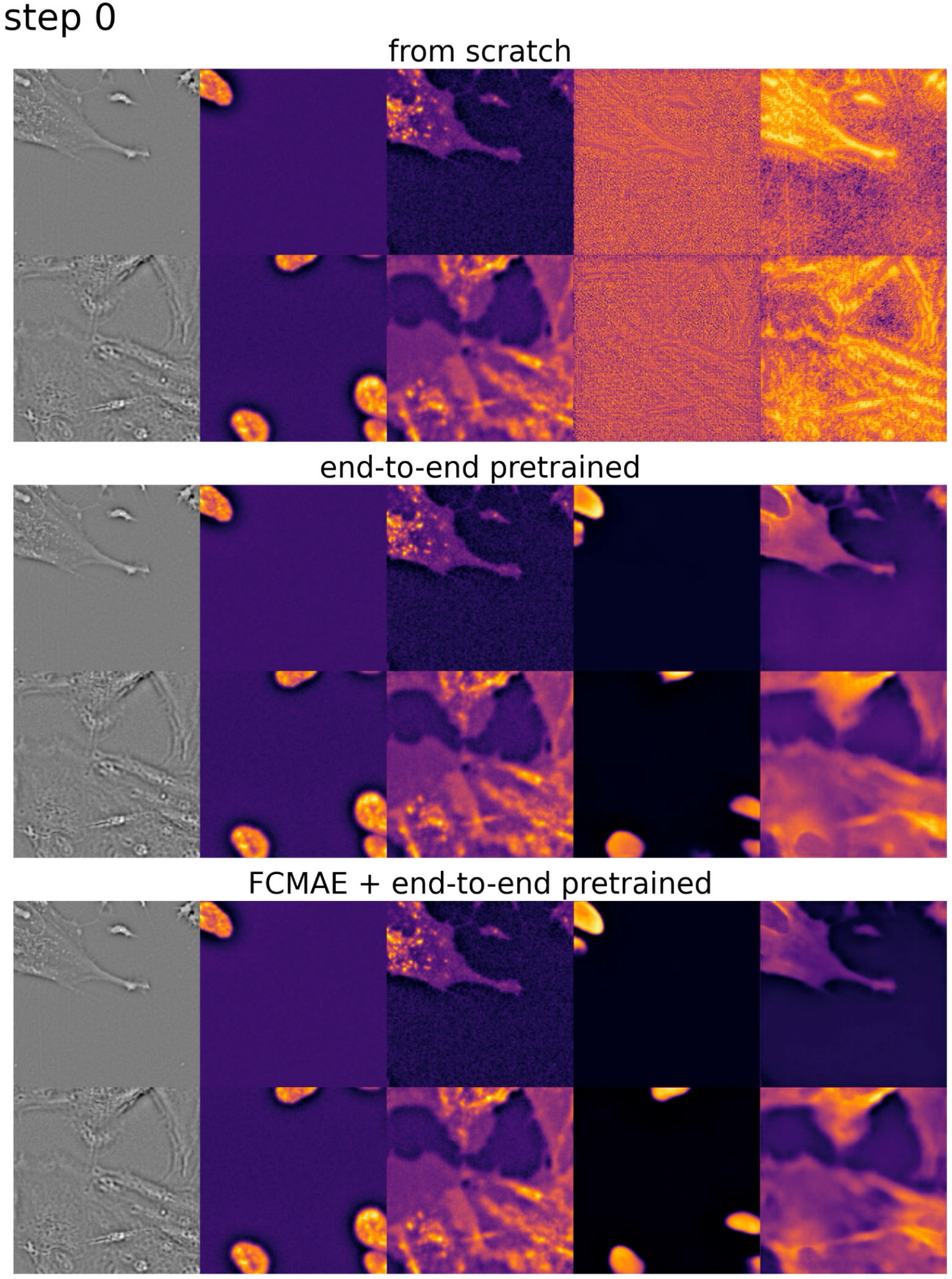
Validation VSCyto2D predictions during fine-tuning on BJ-5ta cells. Each step is 4 training epochs. Left to right: Phase input patch, nuclei fluorescence (Hoechst), membrane fluorescence (CellMask), nuclei prediction, membrane prediction. Pre-trained models start to produce correct predictions faster. Each image patch is 83.2 µm by 83.2 µm (256 pixels by 256 pixels).

## References

1. H. Kobayashi, K. C. Cheveralls, M. D. Leonetti, L. A. Royer, Self-supervised deep learning encodes high-resolution features of protein subcellular localization. Nat. Methods 19, 995–1003 (2022).

2. M.-A. Bray, et al., Cell Painting, a high-content image-based assay for morphological profiling using multiplexed fluorescent dyes. Nat. Protoc. 11, 1757–1774 (2016).

3. Z. Wu, et al., DynaMorph: self-supervised learning of morphodynamic states of live cells. Mol. Biol. Cell 33, ar59 (2022).

4. J. Burgess, et al., Orientation-invariant autoencoders learn robust representations for shape profiling of cells and organelles. Nat. Commun. 15, 1022 (2024).

5. M. P. Viana, et al., Integrated intracellular organization and its variations in human iPS cells. Nature 613, 345–354 (2023).

6. C. J. Soelistyo, G. Vallardi, G. Charras, A. R. Lowe, Learning biophysical determinants of cell fate with deep neural networks. Nat. Mach. Intell. 4, 636–644 (2022).

7. A. M. Valm, et al., Applying systems-level spectral imaging and analysis to reveal the organelle interactome. Nature 546, 162–167 (2017).

8. A. Kumar, et al., Multispectral live-cell imaging with uncompromised spatiotemporal resolution. [Preprint] (2024). Available at: https://www.biorxiv.org/content/10.1101/2024.06.12.597784v1 [Accessed 30 July 2024].

9. A. Jacobo, A. Dasgupta, A. Erzberger, K. Siletti, A. J. Hudspeth, Notch-Mediated Determination of Hair-Bundle Polarity in Mechanosensory Hair Cells of the Zebrafish Lateral Line. Curr. Biol. 29, 3579–3587.e7 (2019).

10. M. N. Hewitt, I. A. Cruz, D. W. Raible, Data-Driven 3D Shape Analysis Reveals Cell Shape-Fate Relationships in Zebrafish Lateral Line Neuromasts. [Preprint] (2023). Available at: https://www.biorxiv.org/content/10.1101/2023.08.09.552694v1 [Accessed 26 May 2024].

11. S.-M. Guo, et al., Revealing architectural order with quantitative label-free imaging and deep learning. eLife 9, e55502 (2020).

12. I. E. Ivanov, et al., Mantis: high-throughput 4D imaging and analysis of the molecular and physical architecture of cells. [Preprint] (2023). Available at: https://www.biorxiv.org/content/10.1101/2023.12.19.572435v1 [Accessed 5 January 2024].

13. I. E. Ivanov, et al., Correlative imaging of the spatio-angular dynamics of biological systems with multimodal instant polarization microscope. Biomed. Opt. Express 13, 3102–3119 (2022).

14. Y. Park, C. Depeursinge, G. Popescu, Quantitative phase imaging in biomedicine. Nat. Photonics 12, 578 (2018).

15. R. Horstmeyer, J. Chung, X. Ou, G. Zheng, C. Yang, Diffraction tomography with Fourier ptychography. Optica 3, 827–835 (2016).

16. O. Liba, et al., Speckle-modulating optical coherence tomography in living mice and humans. Nat. Commun. 8, 15845 (2017).

17. L.-H. Yeh, et al., Permittivity tensor imaging: modular label-free imaging of 3D dry mass and 3D orientation at high resolution. Nat. Methods 1–18 (2024). 10.1038/s41592-024-02291-w.

18. I. E. Ivanov, et al., Mantis: high-throughput 4D imaging and analysis of the molecular and physical architecture of cells. *PNAS Nexus* In Press (2024).

19. J. Ando, A. F. Palonpon, M. Sodeoka, K. Fujita, High-speed Raman imaging of cellular processes. Curr. Opin. Chem. Biol. 33, 16–24 (2016).

20. K. Klein, et al., Label-Free Live-Cell Imaging with Confocal Raman Microscopy. Biophys. J. 102, 360–368 (2012).

21. E. M. Christiansen, et al., In silico labeling: Predicting fluorescent labels in unlabeled images. Cell 173, 792–803.e19 (2018).

22. C. Ounkomol, S. Seshamani, M. M. Maleckar, F. Collman, G. R. Johnson, Label-free prediction of three-dimensional fluorescence images from transmitted-light microscopy. Nat. Methods 15, 917 (2018).

23. J. Park, et al., Artificial intelligence-enabled quantitative phase imaging methods for life sciences. Nat. Methods 20, 1645–1660 (2023).

24. L. Kreiss, et al., Digital staining in optical microscopy using deep learning - a review. PhotoniX 4, 34 (2023).

25. Y. Winetraub, et al., Noninvasive virtual biopsy using micro-registered optical coherence tomography (OCT) in human subjects. Sci. Adv. 10, eadi5794 (2024).

26. B. Bai, et al., Deep learning-enabled virtual histological staining of biological samples. Light Sci. Appl. 12, 57 (2023).

27. Y. Li, et al., Virtual histological staining of unlabeled autopsy tissue. Nat. Commun. 15, 1684 (2024).

28. D. Petersen, et al., Virtual staining of colon cancer tissue by label-free Raman micro-spectroscopy. Analyst 142, 1207–1215 (2017).

29. N. Elmalam, L. Ben Nedava, A. Zaritsky, In silico labeling in cell biology: Potential and limitations. Curr. Opin. Cell Biol. 89, 102378 (2024).

30. M. Pachitariu, C. Stringer, Cellpose 2.0: how to train your own model. Nat. Methods 19, 1634–1641 (2022).

31. C. Stringer, M. Pachitariu, Cellpose3: one-click image restoration for improved cellular segmentation. [Preprint] (2024). Available at: https://www.biorxiv.org/content/10.1101/2024.02.10.579780v2 [Accessed 7 April 2024].

32. A. Archit, et al., Segment Anything for Microscopy. [Preprint] (2023). Available at: https://www.biorxiv.org/content/10.1101/2023.08.21.554208v1 [Accessed 7 April 2024].

33. J. Ma, et al., The multimodality cell segmentation challenge: toward universal solutions. Nat. Methods 1–11 (2024). 10.1038/s41592-024-02233-6.

34. Z. Liu, et al., VisCy: computer vision models for single-cell phenotyping. (2023). Deposited 19 December 2023.

35. J. M. Soto, J. A. Rodrigo, T. Alieva, Label-free quantitative 3D tomographic imaging for partially coherent light microscopy. Opt. Express 25, 15699–15712 (2017).

36. T. Chandler, L.-H. Yeh, I. Ivanov, C. Foltz, S. Mehta, waveorder. (2023). Deposited February 2023.

37. C. Stringer, M. Pachitariu, Transformers do not outperform Cellpose. [Preprint] (2024). Available at: https://www.biorxiv.org/content/10.1101/2024.04.06.587952v1 [Accessed 7 April 2024].

38. S. L. Smith, A. Brock, L. Berrada, S. De, ConvNets Match Vision Transformers at Scale. [Preprint] (2023). Available at: http://arxiv.org/abs/2310.16764 [Accessed 16 May 2024].

39. T. Falk, et al., U-Net: deep learning for cell counting, detection, and morphometry. Nat. Methods 16, 67–70 (2019).

40. S. Woo, et al., ConvNeXt V2: Co-Designing and Scaling ConvNets With Masked Autoencoders in (2023), pp. 16133–16142.

41. K. Tian, et al., Designing BERT for Convolutional Networks: Sparse and Hierarchical Masked Modeling. [Preprint] (2023). Available at: http://arxiv.org/abs/2301.03580 [Accessed 26 May 2024].

42. C. Edlund, et al., LIVECell—A large-scale dataset for label-free live cell segmentation. Nat. Methods 18, 1038–1045 (2021).

43. T. Chen, S. Kornblith, M. Norouzi, G. Hinton, A Simple Framework for Contrastive Learning of Visual Representations in *Proceedings of the 37th International Conference on Machine Learning*, (PMLR, 2020), pp. 1597–1607.

44. M. J. Cardoso, et al., MONAI: An open-source framework for deep learning in healthcare. (2022). 10.48550/arXiv.2211.02701.

45. N. H. Cho, et al., OpenCell: Endogenous tagging for the cartography of human cellular organization. Science 375, eabi6983 (2022).

46. R. Tian, et al., CRISPR Interference-Based Platform for Multimodal Genetic Screens in Human iPSC-Derived Neurons. Neuron 104, 239–255.e12 (2019).

47. J. Peloggia, et al., Adaptive cell invasion maintains lateral line organ homeostasis in response to environmental changes. Dev. Cell 56, 1296–1312.e7 (2021).

48. J. Moore, et al., OME-Zarr: a cloud-optimized bioimaging file format with international community support. Histochem. Cell Biol. 160, 223–251 (2023).

49. Z. Liu, et al., iohub. (2024). Deposited February 2024.

50. T. Chandler, et al., recOrder. (2022). Deposited 23 August 2022.

51. I. E. Ivanov, E. Hirata-Miyasaki, T. Chandler, S. B. Mehta, czbiohub-sf/shrimPy. (2023). Deposited 19 December 2023.

52. H. Zhao, O. Gallo, I. Frosio, J. Kautz, Loss Functions for Neural Networks for Image Processing. [Preprint] (2018). Available at: http://arxiv.org/abs/1511.08861 [Accessed 30 August 2023].

53. Z. Wang, E. P. Simoncelli, A. C. Bovik, Multiscale structural similarity for image quality assessment in The Thirty-Seventh Asilomar Conference on Signals, Systems & Computers, 2003, (IEEE, 2003), pp. 1398–1402.

54. S. van der Walt, et al., scikit-image: image processing in Python. PeerJ 2, e453 (2014).

55. E. Meijering, et al., Design and validation of a tool for neurite tracing and analysis in fluorescence microscopy images. Cytometry A **58A**, 167–176 (2004).

56. N. Otsu, A Threshold Selection Method from Gray-Level Histograms. IEEE Trans. Syst. Man Cybern. 9, 62–66 (1979).

57. T. Y. Zhang, C. Y. Suen, A fast parallel algorithm for thinning digital patterns. Commun. ACM 27, 236–239 (1984).

58. A. Paszke, et al., PyTorch: An Imperative Style, High-Performance Deep Learning Library in Advances in Neural Information Processing Systems 32, H. Wallach, et al., Eds. (Curran Associates, Inc., 2019), pp. 8024–8035.

59. W. Falcon, The PyTorch Lightning team, PyTorch Lightning. (2019). 10.5281/zenodo.3828935. Deposited March 2019.

60. huggingface/pytorch-image-models. (2024). Deposited 19 May 2024.

